# Genome architecture shapes the evolutionary origins of redundant enhancers in fly and mouse

**DOI:** 10.64898/2026.06.26.734851

**Authors:** Jillian Ness, Brendan Kosztyo, Zeba Wunderlich

## Abstract

Shadow enhancers are groups of DNA regulatory elements that control the same target gene and drive overlapping expression patterns. Large-scale surveys have found shadow enhancers control most developmental genes in animal genomes. The way in which shadow enhancers arise and how they subsequently evolve may further illuminate their regulatory logic and mechanisms of action. To investigate the evolutionary origins of shadow enhancers, we searched for sequence signatures of three birth mechanisms: duplication of existing enhancers, transposable element (TE) co-option, and TE-mediated splitting of ancestral regulatory elements in the *Drosophila melanogaster* and mouse genomes. Using 420 fly shadow enhancer sets and 9,051 mouse shadow enhancer sets, we found detectable duplication evidence in 18.3% of fly shadow enhancer sets and 33.9% of mouse sets. Duplication signatures were more frequent in larger shadow sets, suggesting that repeated duplication can expand regulatory landscapes. TE-derived enhancers were present in both species but were not enriched in shadow enhancers relative to single enhancers, suggesting that TE co-option contributes to enhancer evolution generally rather than preferentially generating redundant enhancer architectures. Finally, TE-mediated enhancer splitting was rare in both genomes. These results indicate that shadow enhancer birth is mechanistically heterogeneous, reflecting a mixture of duplication, TE co-option, and other mechanisms, whose contributions are shaped by genome architecture and evolutionary time. Therefore, we find that overlapping regulatory functions can arise through multiple evolutionary routes and that birth mechanisms can influence, but do not strictly determine, the regulatory logic of the resulting enhancer set.

## Introduction

Normal development relies on precise spatiotemporal control of gene expression, and these expression programs are largely encoded in enhancers. Shadow enhancers are a class of enhancers that regulate the same target gene and drive partially or fully overlapping expression patterns^1–3^. These enhancer architectures are widespread across organisms and are particularly common in developmental gene loci^3–5^. Functionally, shadow enhancers can promote phenotypic robustness by buffering genetic, environmental, and molecular perturbations, including transcriptional noise^5–12^. Although shadow enhancers are now recognized as widespread and functionally important, it remains unclear how multi-enhancer architectures arise in the genome. Understanding their origins can help clarify how enhancer redundancy reflects both the mechanisms that generate new regulatory elements and the selective pressures that maintain or refine them.

Several evolutionary routes can generate new enhancers, including duplication of existing enhancers, transposable element (TE) co-option, TE-mediated splitting of ancestral regulatory elements, co-option of unrelated regulatory elements, and de novo gain or reorganization of transcription factor binding sites (TFBS)^3,13^. Duplicated regulatory elements have been described at several loci, supporting the idea that noncoding duplication can contribute to overlapping enhancer architectures^14–17^. TE-derived sequences have contributed to lineage- and stage-specific regulatory elements in mammals and to mammalian shadow enhancer sets^18–21^, while de novo enhancer activity has been observed in *Drosophila*, random sequence libraries, and recently evolved mammalian regulatory elements^22–24^. Because these routes begin from different sequence substrates, they make distinct predictions about shadow enhancer architecture. Duplication can create highly similar enhancer copies with initially shared TFBS content, whereas TE-mediated splitting may separate complementary fragments of an ancestral regulatory grammar. TE-derived and de novo shadow enhancers are expected to show weaker similarity to pre-existing enhancers and may achieve overlapping expression through TFBS gain, loss, or turnover. Thus, identifying birth routes can help explain why some shadow enhancers share regulatory logic, whereas others achieve overlapping expression through distinct sequence features.

Here, we investigated the evolutionary origins of shadow enhancers in the *Drosophila melanogaster* and mouse genomes by searching for sequence signatures of three birth mechanisms: duplication of existing enhancers, transposable element (TE) co-option, and TE-mediated splitting of ancestral regulatory elements. To characterize these routes of birth, we used 420 fly shadow enhancer sets comprising 1,122 enhancers, including developmental enhancers identified using in vivo reporters and mesoderm enhancers identified using bioinformatic analysis of TF binding and gene expression data^4,25^. We also used 9,051 mouse shadow enhancer sets comprising 19,166 enhancers active in limb, heart, and forebrain development and identified via bioinformatic analysis of histone H3K27ac mark, gene expression, and 3D chromatin conformation data^5^. This comparison allowed us to test whether shadow enhancer birth mechanisms vary between genomes that differ in size, repeat content, and evolutionary history.

We find that shadow enhancer birth is mechanistically heterogeneous. Duplication signatures are detectable in both species and are more common in larger shadow sets, suggesting that duplication can contribute to the expansion of existing regulatory landscapes. TE-derived enhancers occur in a minority of shadow sets, are more frequent in the TE-rich mouse genome than fly, and are not enriched in shadow enhancers relative to single enhancers. Finally, TE-mediated splitting is rare in both genomes, suggesting that many shadow enhancer sets may arise through de novo enhancer emergence, regulatory co-option, ancient duplication, or older TE-associated events whose sequence signatures are no longer detectable. Below, we begin by considering each shadow enhancer birth route in the *Drosophila melanogaster* data set and then analyze the mouse data set.

## Results

### Putative shadow enhancer duplications were identified using enhancer and flanking sequence comparisons

Given the shared function of sets of shadow enhancers, a natural hypothesis for their emergence is duplication of an existing enhancer. We analyzed 1,122 *Drosophila* developmental shadow enhancers across 420 shadow sets (median = 2 enhancers/set; range = 2–13) and screened each set for two duplication signatures: enhancer-body similarity and flanking-sequence homology (Figure 1A). Enhancer-body homology was identified using pairwise BLAST alignments between enhancers within each set, with hits called using an empirically calibrated cutoff (E ≤ 10^-4^; FPR ≈ 5%; Methods). This enhancer-hit cohort included 81 enhancer pairs across 56 shadow sets, corresponding to 13.3% of fly shadow sets (Figure 1B,C). Aligned regions had high sequence identity (median = 93.3%) but were typically short, with a median length of 18 bp (Supplementary Figure 1, Supplementary Table 1). These short but highly conserved sequences are consistent with studies of the evolution of homologous enhancers between species, where similarly short, highly conserved sequences are detectable in functionally conserved enhancers^26–28^.

**Figure 1.**
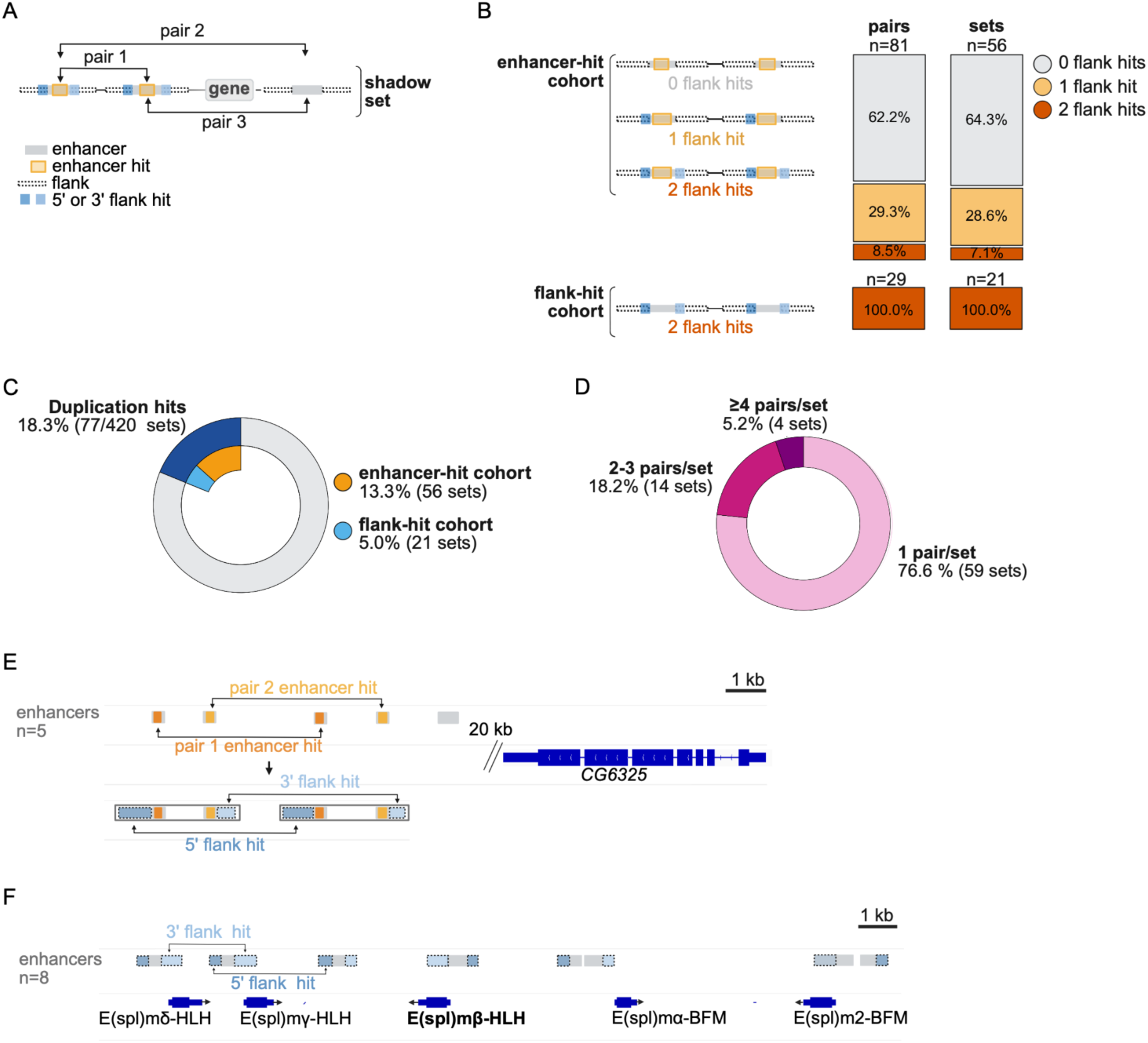
Enhancer-body and flanking sequence homology identify putative duplications in fly shadow enhancer sets. **(A)** Schematic defining duplication cohorts. Sequence alignments were performed on shadow enhancers pairs and their flanking regions. The enhancer-hit cohort is composed of pairs with a significant enhancer-body alignment (BLAST, E ≤ 10^-4^), optionally supported by 5′/3′ flank homology. The flank-only cohort is composed of pairs with paired 5′ and 3′ flank homology but no enhancer-body hit. **(B)** For enhancer-hit pairs, flanking regions were evaluated for additional evidence of duplication. Stacked bars show the fraction of enhancer-hit *pairs* (n = 81) or enhancer-hit *sets* (n = 56) with no detectable flank homology (grey), one flank hit (orange), or paired 5′/3′ flank hits (red). The flank-hit cohort includes enhancer pairs lacking enhancer-body similarity but recovered by paired 5′ and 3′ flank homology (red, n = 29 pairs across 21 sets). **(C)** A donut plot of the total fraction of shadow enhancer sets with detectable duplication evidence. Duplication hits (blue) are partitioned into enhancer-hit sets (orange) and flank-hit sets (light blue). **(D)** Percentage of duplication-hit shadow sets grouped by the number of hit pairs per set: 1 hit pair/set (pink), 2–3 hit pairs/set (dark pink), or ≥4 hit pairs/set (magenta). **(E)** The *CG6325* locus is regulated by five shadow enhancers and contains two duplicated enhancer pairs that may have arisen from a tandem duplication of two enhancers. Enhancers are shown in gray, duplicated enhancer-body regions in orange, and homologous 5′ and 3′ flanks in blue/light blue. (**F**) The *E(spl)mbeta* shadow set duplications can be identified using flank analysis. The eight shadow enhancers are shown in gray, and homologous 5′ and 3′ flanks in blue/light blue.

We next considered two caveats of the BLAST-based approach: false positives from convergent TFBS clustering and false negatives from enhancer sequence turnover. To assess convergence, we scanned fly BLAST hits for overlap with JASPAR *D. melanogaster* TF binding motifs. Only ∼6% of hits (5/88) were ≥80% covered by predicted TFBSs, suggesting that most BLAST hits are more consistent with shared ancestry than convergent motif clustering (Supplementary Figure 2). To assess sensitivity of our BLAST approach, we tested recovery of functionally conserved homologous enhancers^22^. BLAST recovered ∼85% of homologous sequences across ∼50–60 Myr of divergence, but only ∼5% across ∼250 Myr (Supplementary Figure 3). We therefore interpret enhancer-hit calls as recent-to-intermediate duplication signatures, while recognizing that older duplications are likely underestimated.

Because segmental duplication mechanisms copy contiguous DNA blocks, enhancers can be duplicated with their surrounding sequence, including their nearby genes^29^. Therefore, we reasoned that we could analyze sequences flanking enhancers both to further validate our BLAST-based duplication calls and potentially uncover other duplication events. We assayed homology in 5′ and 3′ flanks for all shadow enhancer pairs. Among enhancer-hit pairs, flank conservation was frequent, with 29.3% of pairs showing a single flank hit, 8.5% showing paired 5′+3′ flank hits, and 62.2% had no detectable flank homology (Figure 1B). Therefore, flanks can provide independent evidence for duplication for some shadow enhancer pairs.

We then tested whether paired flank homology alone—in the absence of enhancer-body similarity—can uncover additional events.This identified a flank-only cohort composed of 29 shadow enhancer pairs in 21 additional sets, in which there is paired 5′ and 3′ flank homology despite no significant enhancer-body hit (Figure 1B). Together, the two approaches yield 18.3% of shadow sets with duplication hits, defined by two non-overlapping duplication cohorts: an enhancer-hit cohort (13.3%) and a flank-only cohort (5%) (Figure 1C). These results suggest that matching flanks can expose duplicated enhancers even when the enhancer cores have diverged over evolutionary time.

Because enhancer sets may undergo repeated duplication, we hypothesized that duplication signatures may extend beyond single enhancer pairs. Among the 77 fly shadow sets with duplications, 18 sets (24%) contained multiple duplicated pairs (Figure 1D). One example is the mesoderm-active *CG6345* locus, which contains five shadow enhancers, including two enhancer pairs with strong signatures of recent duplication (Figure 1E). Both pairs share high enhancer-body similarity and homologous 5′ and 3′ flanking sequence, consistent with a duplication event that encompassed multiple enhancers and surrounding sequence (Supplementary Figure 4). This example illustrates how local duplication can rapidly enlarge a shadow enhancer set and how flanking homology can help localize duplication boundaries.

The eight shadow enhancers in the duplicated *E(spl)m* multi-gene loci demonstrate the utility of the flanking sequence analysis (Figure 1F). The *E(spl)m* expansion predates schizophoran diversification and the locus shows strong conservation of gene order and regulatory organization^30,31^. These duplicated mesoderm enhancers are annotated as regulating *E(spl)mβ*, because it is the only local paralog with mesoderm expression, though it is plausible the enhancers may act on multiple *E(spl)m* promoters in other tissue and developmental contexts. Among the eight shadow enhancers, enhancer-body similarity is weak, but flanking sequence analysis reveals shared 5′ and 3′ sequences across the set (Figure 1F). This pattern is consistent with enhancers being carried through serial gene duplications, and demonstrates that residual flank conservation can reveal duplication events when enhancer-body similarity is weak.

### Larger fly shadow enhancer sets are more likely to arise from duplications than smaller sets

A number of shadow sets have evidence of multiple duplication events, and we hypothesized that duplication can rapidly enlarge shadow enhancer set size. We calculated the fraction of enhancers participating in duplication-hit pairs within each shadow set size bin (Figure 2A). This fraction increased with set size, from 9.3% in two-enhancer sets to 16.8% in three-enhancer sets and 24.5% in sets with at least four enhancers (Supplementary Figure 5). Each bin contained significantly more duplication-hit enhancers than expected from size-matched randomized controls (P < 10^-6^; Supplementary Figure 6). Larger set-size bins had significantly higher duplication-hit proportions than smaller bins (Supplementary Figure 5), supporting a model in which larger shadow sets are more likely to have expanded via duplication.

**Figure 2.**
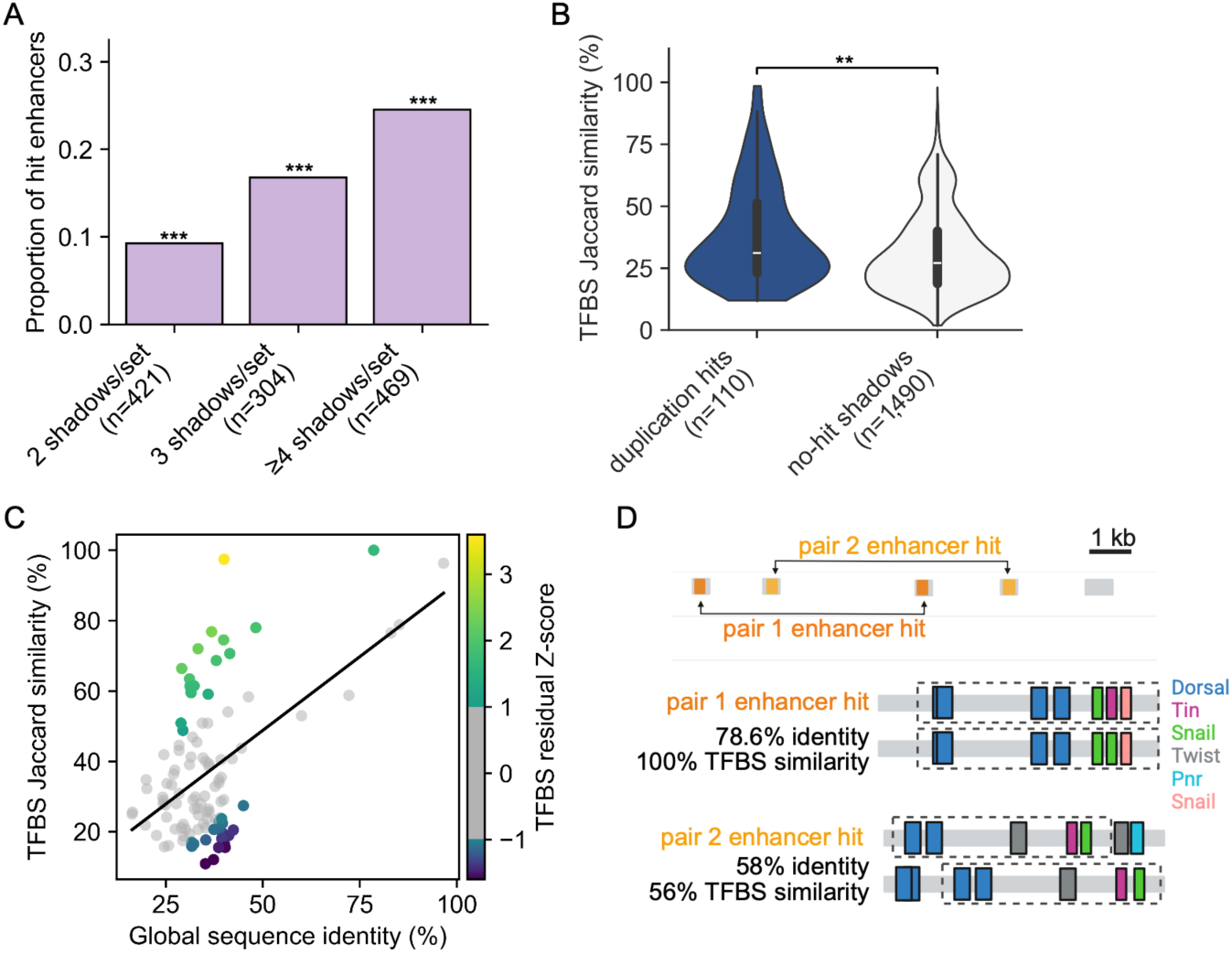
Fly shadow enhancer duplications are enriched in larger sets and show variable TFBS conservation. **(A)** Proportion of hit enhancers grouped by shadow set size: 2 shadows/set (n = 421 enhancers), 3 shadows/set (n = 304 enhancers), or ≥4 shadows/set (n = 469 enhancers), where (n) indicates the total number of enhancers in each set-size bin. Asterisks indicate significance relative to size-matched randomized control distribution (*P < 0.01, **P < 0.001, ***P < 10^-6^). **(B)** TFBS Jaccard similarity for duplication-hit pairs (blue; n = 110 pairs) compared with no-hit (grey; n = 1,490 pairs) shadow pairs. Significance was assessed using a Mann–Whitney U test (*P < 0.01, **P < 0.001, ***P < 10^-6^). **(C)** Global sequence identity versus TFBS Jaccard similarity for duplication-hit pairs. The black line shows a linear regression fit. Point color indicates TFBS residual z-score, highlighting pairs with higher- or lower-than-expected TFBS similarity given their sequence identity. **(D)** The *CG6325* locus (as in Figure 1E) is shown with the TFBS and global sequence similarly of putative duplicated pair 1 (dark orange) and pair 2 (light orange). Diagrams show the TFBS content (colored rectangles) for each pair of enhancers, and the boxed regions denote the region of the BLAST hit. (*P < .01, **P < .001 and ***P < 10^-6^).

### Duplicated fly shadow enhancers may retain or diverge in their TFBS content

After an enhancer duplicates, there are several possible evolutionary trajectories. The resulting shadow enhancers may retain overlapping function through conserved TFBS content, or through sequence turnover that preserves activity while altering TFBS composition. To assess the degree of regulatory architecture conservation between duplicated shadow enhancers, we quantified TFBS similarity using a pairwise similarity metric for 14 mesoderm transcription factor motifs (Methods). Duplication-hit pairs showed significantly higher TFBS similarity than no-hit shadow pairs (P < 10^-3^, Figure 2B), consistent with their shared evolutionary origin and partial retention of regulatory logic after duplication. But, the broad overlap between distributions suggests heterogeneous evolutionary outcomes: some duplicated pairs retain similar motif content, whereas others diverge in TFBS composition.

To interpret TFBS similarity in the context of sequence divergence, we compared TFBS similarity with global enhancer sequence identity (Figure 2C). Most duplicated pairs fell near the fitted relationship, but some showed higher-than-expected TFBS similarity, consistent with retention of binding-site architecture despite broader sequence divergence. Others showed lower-than-expected TFBS similarity, consistent with regulatory rewiring despite moderate sequence conservation.

To illustrate these post-duplication trajectories, we again examined the *CG6325* shadow enhancer set, which contains five shadow enhancers and two putative duplicated enhancer pairs (Figure 2D). Both pairs show substantial enhancer-body sequence similarity, consistent with recent local duplication, but they differ in the extent to which predicted TFBS organization is retained. One pair has complete TFBS overlap, whereas the second pair shows lower TFBS similarity despite detectable sequence homology. Thus, *CG6325* provides an example of how enhancer duplication can expand a shadow enhancer set while allowing the duplicated elements to retain or partially diverge in regulatory logic after birth.

Overall, while DNA duplication is a plausible route of shadow enhancer genesis, our analysis suggests that the mechanism does not explain the origins of the majority of shadow enhancers in the *Drosophila* genome. While we acknowledge caveats to the analysis, e.g. the inability to detect very ancient duplication events, we propose other routes of shadow enhancer birth may also be prevalent.

### A small fraction of fly shadow enhancers are TE-derived

TEs are a potential source of new enhancer activity because their insertions can carry new sequence, sometimes including TFBS, into existing gene regulatory landscapes. Their co-option into enhancers can arise from these pre-existing TFBS or accumulation of TFBS after insertion. To assess the prevalence of TE-derived shadow enhancers, we annotated TEs in the *Drosophila* genome with curated and de novo annotations, producing a TE set spanning ∼21% of the genome, consistent with recent TE occupancy estimates^32^. We classified enhancers overlapping TE sequence by at least 20% as TE-derived, or TE+, allowing us to identify shadow enhancer sets in which members bear TE ancestry (Figure 3A).

**Figure 3.**
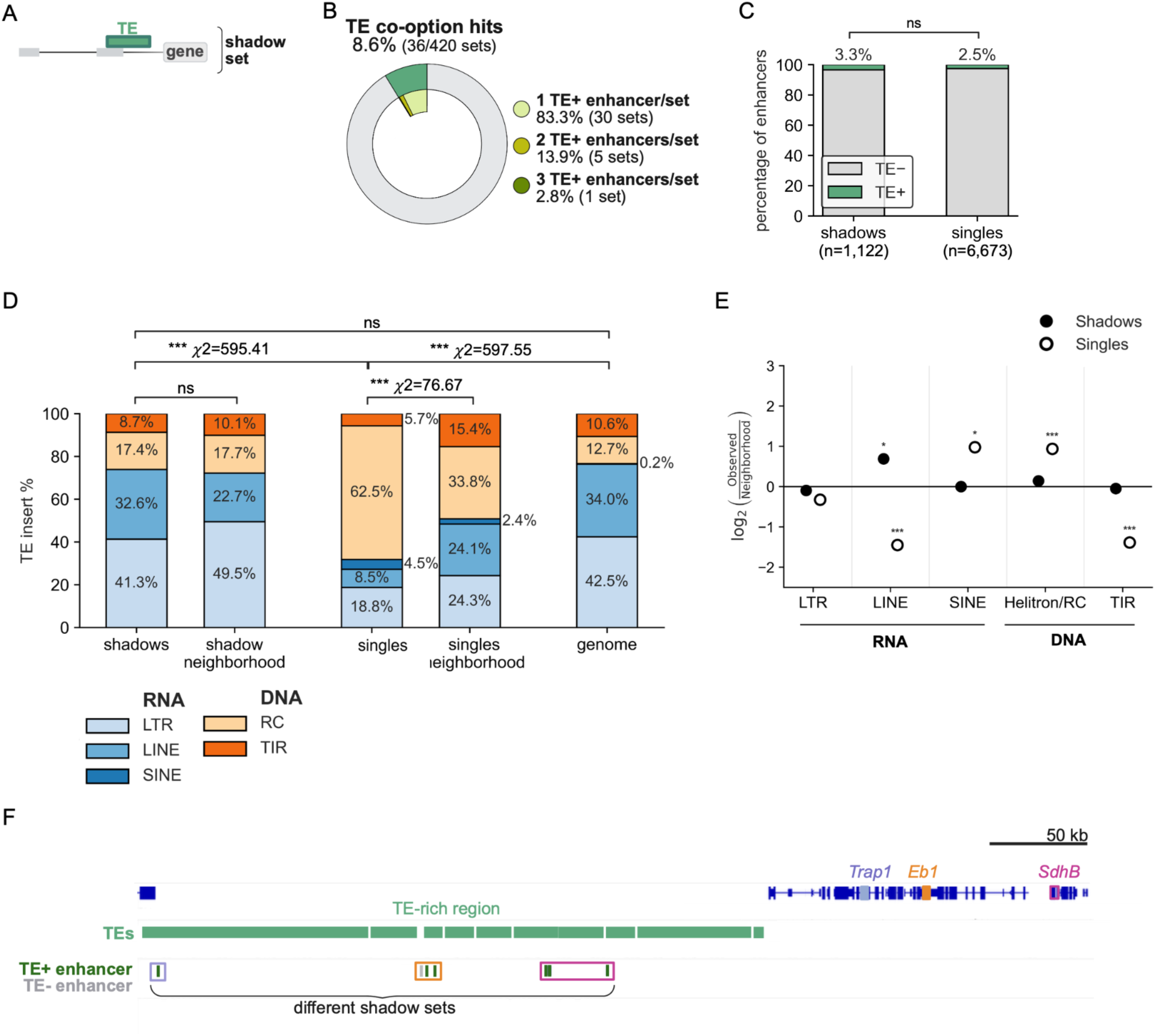
Fly TE co-option is rare, partly reflects local TE availability, and is not predicted by TE-class motif content. **A)** Schematic of a TE-derived (TE+) enhancer that functions as a member of a shadow enhancer set. Enhancers were classified as TE+ if annotated TE sequence overlapped at least 20% of the enhancer length. (**B)** The percentage of fly shadow enhancer sets that contain at least one TE-derived enhancer, termed TE+ sets, or no TE-derived enhancers, termed TE− sets. The inner donut shows the proportion of TE+ sets containing one, two, or three TE+ enhancers. **(C**) Percent of TE+ enhancers in the shadow data set (n = 1,122) or the single enhancer dataset (n = 6,673). ns, not significant (two-proportion z-test, P = 0.31). **(D)** Stacked bars show the TE class composition of TE+ shadow enhancers, TE+ single enhancers, their matched local neighborhoods, and the genome. Local neighborhoods were defined as the 50 kb upstream and downstream regions surrounding each TE+ enhancer. TE classes are grouped as RNA-derived elements, including LTR, LINE, and SINE elements, and DNA-derived elements, including Helitron/RC and TIR elements. Asterisks indicate significance from chi-square goodness-of-fit tests (ns = not significant, *P < 0.01, **P < 0.001, ***P < 10^-6^). **(E)** Log_2_ enrichment of TE classes among TE+ enhancers relative to their matched local neighborhoods. Filled circles indicate shadow enhancers and open circles indicate single enhancers. Asterisks indicate FDR-adjusted significance for per-class observed-versus-neighborhood enrichment tests (ns = not significant, *P < 0.01, **P < 0.001, ***P < 10^-6^). **(F)** Example of a TE-rich noncoding region containing TE+ enhancers from multiple shadow enhancer sets, including the *Trap1*, *Eb1*, and *SdhB* shadow sets.

Across shadow enhancer sets, 8.6% (n = 36) contain at least one TE+ enhancer. Most TE+ sets contain only one TE+ member (83.3%, n = 30), fewer contain two (13.9%), and only one set contains three (2.8%) (Figure 3B). Larger shadow enhancer sets did not have more TE+ enhancers than smaller ones (Supplementary Figure 7). To interpret whether TE+ enhancers are specifically associated with shadow architectures, we analyzed mesoderm “single enhancers” – enhancers not part of a shadow set – as a baseline. TE-derived sequence is rare in both regulatory architectures but occurs at a slightly elevated rate within shadow enhancers, comprising 3.3% of shadow enhancers and 2.5% of single enhancers (Figure 3C).

If co-opted TE insertions are older or under strong constraint, they should be more likely to be fixed across *D. melanogaster* lines. In contrast, recently inserted, recently co-opted, or weakly constrained TEs should be more often polymorphic and present in only a subset of lines. To determine whether TE+ enhancers are fixed in the *D. melanogaster* genome, we scored TE+ enhancers across *D. melanogaster* strain assemblies^33^, and called the TE as fixed when present in at least 90% of lines (Methods)^33^. Among TE+ shadow enhancers, 73.0% (27/37) were fixed and 27.0% (10/37) were not fixed (Supplementary Table 2). Among TE+ single enhancers, 93.9% (169/180) were fixed and 6.1% (11/180) were not fixed (Supplementary Table 3). Thus, TE+ enhancers in shadow architectures are more often polymorphic than those at single enhancer loci (P = 5 × 10^-4^, Fisher’s exact test), consistent with more recent insert or co-option, weaker constraint, or a combination of factors.

### TE class composition of co-opted fly enhancers broadly follows local genomic TE neighborhood

TE co-option frequencies may vary from class to class, either because of their local prevalence or sequence material. If some TE classes have greater mesodermal proto-enhancer potential, they could contribute disproportionately to TE-derived enhancers. TE-class composition differed strongly between TE+ shadow and single enhancers: shadow enhancers were enriched for RNA-based elements, especially Long Terminal Repeat retrotransposons (LTRs) and Long Interspersed Nuclear Elements (LINEs), whereas single enhancers were dominated by DNA transposons, particularly Helitron/Rolling Circle (Helitron/RC) elements (Figure 3D). To determine whether these simply reflect the TE landscape surrounding each enhancer, we compared TE+ enhancers with their local upstream and downstream neighborhoods. Both shadow and single enhancers more closely resembled their local neighborhoods than the genome-wide TE distribution, but each also deviated from neighborhood expectations (Figure 3D,E). Shadow enhancers showed slight LINE enrichment and mild LTR depletion, whereas single enhancers showed stronger enrichment for SINE and Helitron/RC elements and depletion of LTR, LINE, and Terminal Inverted Repeat (TIR) DNA elements (Figure 3E). These trends held when TE composition was measured by overlapping TE base pairs rather than insertion counts (Supplementary Figure 8). We also measured TE class mesoderm TFBS content and compared this to TE+ enhancer TFBS content, but did not find this predicted which TE classes were preferentially co-opted (Supplementary Figure 9). Thus, local TE availability explains part, but not all, of the difference between TE-derived shadow and single enhancers.

Consistent with a role for local TE-rich sequence, we identified noncoding regions containing multiple TE+ shadow enhancer sets. One such region contains TE+ enhancers associated with *Trap1*, *Eb1*, and *SdhB* (Figure 3F). This locus illustrates how TE-dense genomic neighborhoods can provide repeated opportunities for enhancer co-option across different TE classes.

Together, these results suggest that fly TE co-option is shaped partly by local TE availability. Though TE-class TFBS content did not predict TE-co-option, other features, including finer-scale TE family differences, insertion age, chromatin context, and post-insertion sequence evolution, may influence whether a TE-derived sequence becomes incorporated into an enhancer.

### TE-mediated splitting is rare in fly shadow enhancer sets

TE insertions can alter cis-regulatory architecture not only by donating enhancer sequences but also by physically separating pre-existing regulatory elements. If such insertions occur within an ancestral enhancer, they could fragment into two shadow enhancers (Figure 4A). To test how often TEs splitting may contribute to shadow enhancer birth, we examined the inter-enhancer space within shadow enhancer sets and identified cases in which TE sequence accounts for most of the intervening interval.

**Figure 4.**
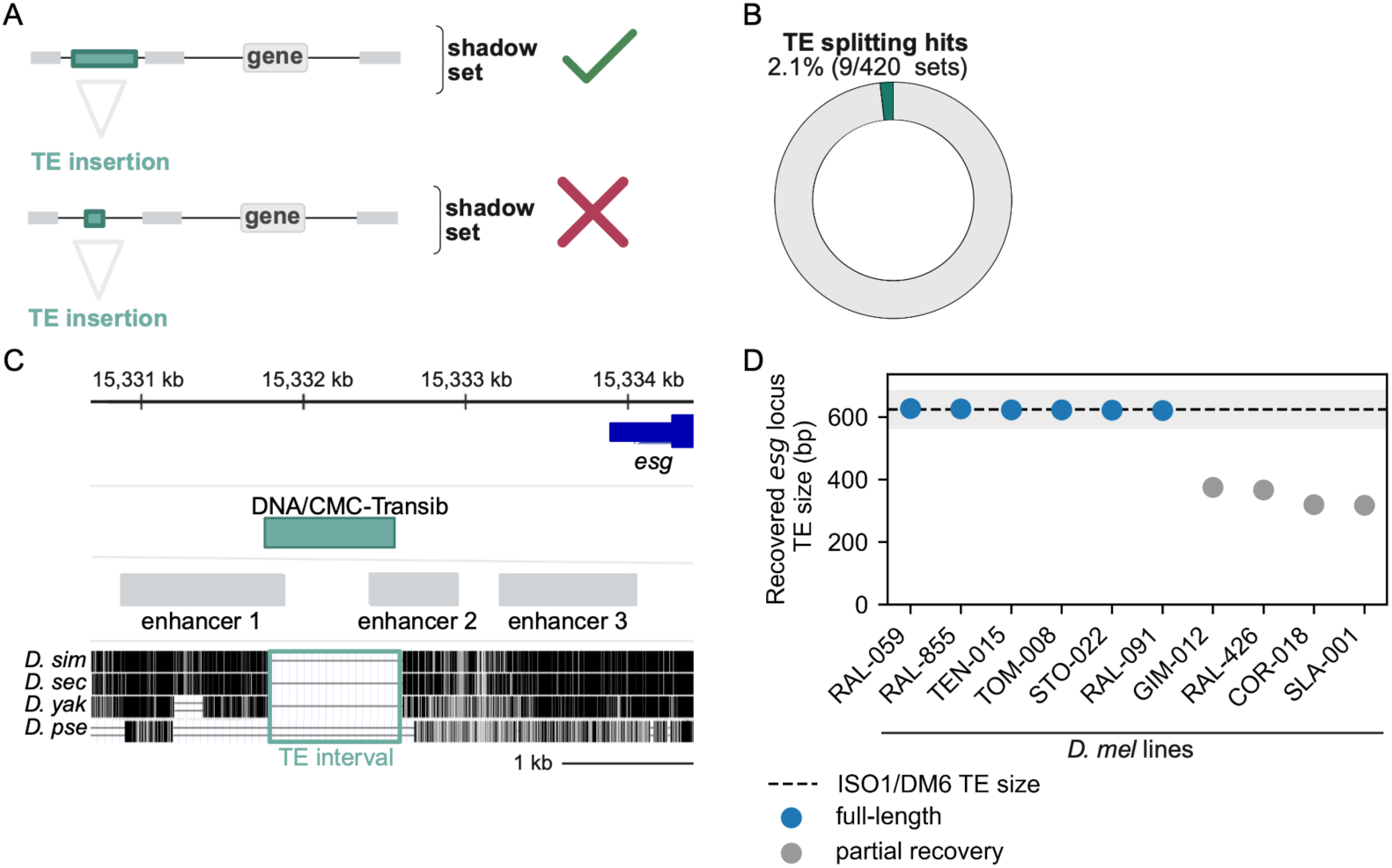
TE-mediated enhancer splitting is rare in fly shadow enhancer sets. **(A)** Schematic illustrating candidate TE-mediated enhancer splitting events, in which TE sequence occupies most of the intervening space between two shadow enhancers, versus apparent non-splitting events, in which the TE appears to lie between two already separate enhancers. **(B)** Fraction of fly shadow enhancer sets containing at least one candidate TE-mediated splitting event. We identified 11 potential splitting events in nine shadow enhancer sets. Four splitting events met the 80% TE-coverage criterion, one had <500 bp of non-TE sequence, and six events satisfied both thresholds. **(C)** Example of a putative enhancer splitting event in the *escargot* (*esg*) shadow enhancer set. The early developmental *esg* shadow set contains six enhancers, two of which are separated by an LTR transposon and are consistent with birth from an ancestral enhancer that was split by TE insertion. This insertion is lineage-specific to *D. melanogaster* and is absent from the orthologous region in both the closely-related species *D. simulans* and the more distantly-related species *D. pseudoobscura*. **(D)** Recovered TE lengths at the candidate *escargot* (*esg*) splitting locus across ten long-read *D. melanogaster* strain assemblies. The TE insertion is recovered across *D. melanogaster* lines, although the inferred TE length varies among assemblies.

We first examined the intervening sequence between all shadow enhancer pairs. Although many intervening sequences contained at least one TE, most of the length of these TE+ intervening sequences was still primarily composed of non-TE sequence (Supplementary Figure 10). Therefore, we defined TE-splitting candidates using two criteria: either the TE sequence occupies at least 80% of the intervening sequence or less than 500 bp of non-TE sequence remains within a TE-containing sequence. These thresholds isolate the subset of loci with unusually TE-dominant intervening space (Supplementary Figure 10). Using these criteria, we identified 11 candidate cases across nine shadow enhancer sets (Figure 4B).

As an example, we highlight a TE insertion located 5’ of the *esg* gene, between enhancers 1 and 2 of a six shadow enhancer set (Figure 4C). This TE is fixed across 10 long-read *D. melanogaster* genome assemblies (Figure 4D; Methods)^33^, although the recovered TE sequence varies in length. Multiple sequence alignments suggest that this TE is lineage-specific to *D. melanogaster* (Figure 4C). Overall, these results suggest that TE-mediated enhancer splitting is not a common route for generating shadow enhancers. Our data support a model in which TE insertions more often contribute to the expansion of existing regulatory landscapes, while only a small subset of cases are consistent with TE-driven enhancer splitting.

### Putative shadow enhancer duplications are common in mouse shadow sets

To examine whether the same routes of shadow enhancer birth observed in *Drosophila* also operate in a divergent genome, we extended our analysis to the mouse genome. It provides a useful comparison because its genome differs substantially from *Drosophila* in size, repeat content, and regulatory landscape, allowing us to test whether duplication and TE-associated mechanisms are shared features of shadow enhancer evolution or instead reflect genome-specific processes. In a data set of shadow enhancers active in limb, forebrain, and heart development, there are 19,166 unique shadow enhancers across 9,051 total shadow sets, with a median of 3 shadows per set and a range of 2 to 43 shadows per set^5^.

To identify shadow enhancer birth by duplication, we applied the same two-pronged approach used above, identifying duplication signatures from either enhancer-body sequence similarity or paired flanking sequence homology (Figure 5A, B). Across the mouse shadow enhancer dataset, we identified enhancer-body homology in 31.9% of shadow enhancer sets (Figure 5C, Supplementary Figure 11, Supplementary Table 4). Among enhancer-hit pairs, flank conservation was frequent, with 63.4% showing a single flank hit and 5.1% showing paired 5′+3′ flank hits, while 31.5% had no detectable flank homology (Figure 5B). These results indicate that a majority of mouse enhancer-hit pairs retain homology in their surrounding sequence, consistent with duplication of a larger local interval, while some enhancer-hit pairs only retain detectable similarity within the enhancer body. We also identified a flank-hit cohort composed of 2.0% of shadow enhancer sets (Figure 5C). Together, the two approaches identified duplication hits in 33.9% of mouse shadow enhancer sets, which is substantially higher than the duplication hit rate in flies (18.3% of fly shadow enhancer sets, Figure 1C; P = 1.39 × 10^-10^, two-proportion Z-test).

**Figure 5.**
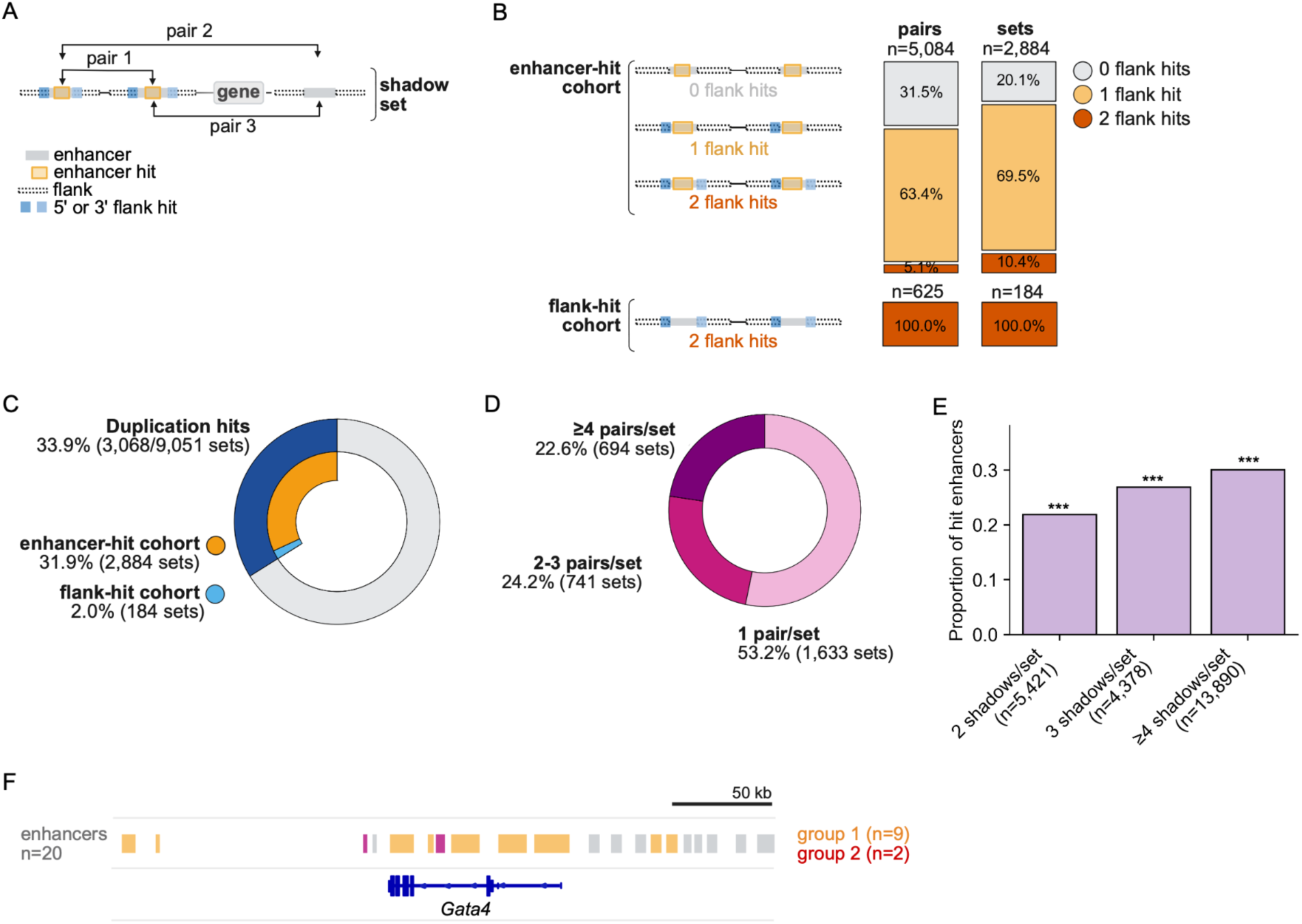
Enhancer-body and flanking-sequence homology identify widespread duplications in mouse shadow enhancer sets. **(A)** Schematic defining mouse duplication cohorts. Sequence alignments were performed on shadow enhancers pairs and their flanking regions. The enhancer-hit cohort is composed of pairs with a significant enhancer-body alignment (BLAST, E ≤ 10^-8^), optionally supported by paired 5′/3′ flank homology. The flank-only cohort is composed of pairs with paired 5′ and 3′ flank homology but no enhancer-body hit. **(B)** Mouse enhancer-hit pairs were first identified by significant enhancer-body similarity. For these pairs, flanking regions were then evaluated for additional evidence of duplication. Stacked bars show the fraction of enhancer-hit *pairs* (n = 5,084) or across enhancer-hit *sets* (n = 2884) with no detectable flank homology (grey), one flank hit (orange), or paired 5′/3′ flank hits (red). The flank-hit cohort is shown separately and includes only enhancer pairs lacking enhancer-body similarity but recovered by paired 5′/3′ flank homology (red, n = 625 pairs across 184 sets). **(C)** The total fraction of shadow enhancer sets with detectable duplication evidence. The total duplication-hit class (blue) is partitioned into two non-overlapping cohorts: enhancer-hit sets (orange), and flank-hit sets (light blue). **(D)** Percentage of duplication-hit shadow sets grouped by the number of hit pairs per set: 1 hit pair/set (pink), 2–3 hit pairs/set (dark pink), or ≥4 hit pairs/set (magenta). **(E)** Proportion of hit enhancers grouped by shadow set size: 2 shadows/set (n = 5,421 enhancers), 3 shadows/set (n = 4,378 enhancers), or ≥4 shadows/set (n = 13,890 enhancers) where (n) indicates the total number of enhancers in each set-size bin. Asterisks indicate significance relative to size-matched randomized control distribution (*P < 0.01, **P < 0.001, ***P < 10^-6^). **(F)** Example of a putative duplication-rich shadow enhancer set at the *Gata4*locus, which contains 20 shadow enhancers. Based on enhancer-body similarity, the locus contains two primary duplication groups with at least 100 bp of shared enhancer homology. Group 1 contains nine enhancers consistent with a series of duplication events, and group 2 contains two enhancers with shared sequence similarity.

Because mouse shadow enhancer sets are larger on average than fly shadow sets, we tested whether mouse sets frequently contain multiple duplicated enhancer pairs and whether duplication-hit enhancers are enriched in larger sets. Among the 3,068 mouse shadow sets with duplication evidence, 53.2% contained one hit pair, 24.2% contained 2–3 hit pairs, and 22.6% contained at least four hit pairs (Figure 5D). The fraction of enhancers participating in duplication-hit pairs increased with set size, from 21.9% in two-enhancer sets to 26.9% in three-enhancer sets and 30.1% in sets with at least four enhancers (Figure 5E; Supplementary Figure 12). Each bin contained significantly more duplication-supported enhancers than expected from size-matched randomized controls (P < 10^-6^) and duplicated enhancers were more common in larger shadow enhancer sets (Supplementary Figure 13; Methods). This indicates that duplication hits are enriched across mouse shadow enhancer sets compared to the fly and are especially frequent in larger sets.

As an example, we highlight the *Gata4* locus, which contains 20 enhancers and multiple putative duplicated enhancer groups (Figure 5F). Several enhancers near *Gata4* have shared sequences consistent with duplication, including a larger group containing nine enhancers and a smaller duplication group containing two enhancers. This locus illustrates how duplication can contribute to expanded shadow enhancer architectures in the mouse genome. Thus, duplication appears to be a shared mechanism of shadow enhancer birth across divergent genomes, but its detectable contribution is greater in the mouse genome than in *Drosophila*.

### TE class co-option patterns are shared between mouse shadow and single enhancers but differ from local TE neighborhoods

We next examined whether TEs also contribute to shadow enhancer birth in the mouse genome through enhancer co-option (Figure 6A). Using the same TE-overlap approach as in the fly genome, we classified enhancers overlapping TE sequence by at least 20% as TE-derived, or TE+. Across mouse shadow enhancer sets, 19.7% contained at least one TE+ enhancer (Figure 6B). Within these TE+ sets, most contained only one TE+ enhancer per set (86.6%), and fewer contained two TE+ enhancers (11.2%) or three TE+ enhancers (2.2%), with no enrichment of TE+ enhancers in larger shadow sets (Supplementary Figure 14). To determine whether TE-derived sequence is specifically associated with shadow enhancer architectures, we compared TE+ enhancer frequency between shadow enhancers and single enhancers and found no difference, with 7.5% of shadow and single enhancers classified as TE+ (Figure 6C). Thus, TE co-option is present in a minority of mouse shadow enhancer sets, is not enriched in shadow enhancers relative to single enhancers, and usually affects only a single enhancer within a set.

**Figure 6.**
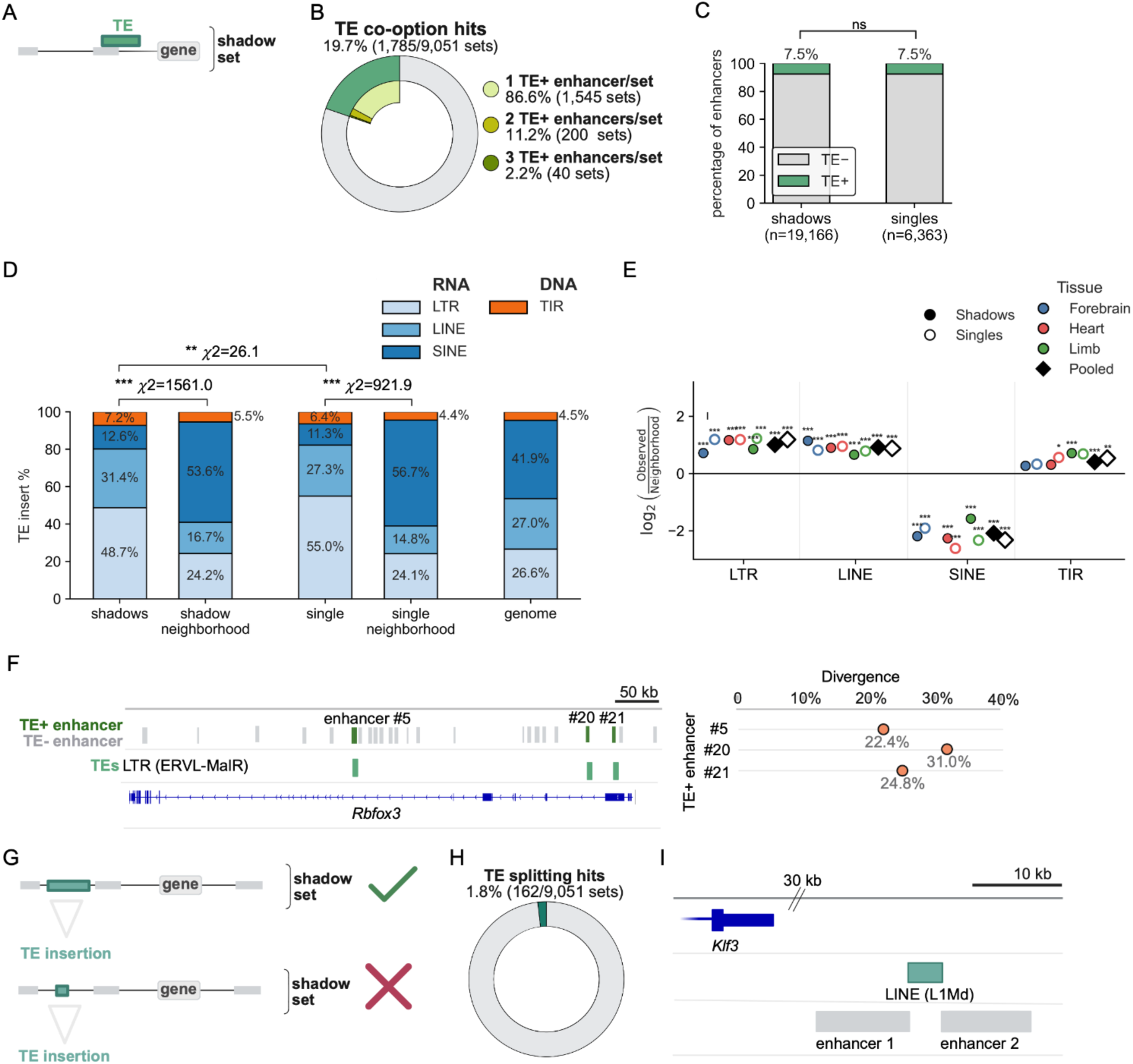
Mouse TE co-option is more common than in fly, reflects TE-class bias, and TE-mediated splitting is rare. **(A)** As in Figure 3A, an enhancer is called TE+ is the TE takes up at least 20% of the enhancer length. **(B)** The percentage of mouse shadow enhancer sets that contain at least one TE-derived enhancer, termed TE+ sets, or no TE-derived enhancers, termed TE− sets. The inner donut shows the proportion of TE+ sets containing one, two, or three TE+ enhancers. **(C)** Percentage of TE+ enhancers among mouse shadow enhancers (n = 19,166) and single enhancers (n = 6,363). ns, not significant (two-proportion Z-test, P = .99) **(D)** Stacked bars show the TE’s of the TE+ shadow enhancers, TE+ single enhancers, their matched local neighborhoods, and the genome. Local neighborhoods were defined as the 50 kb upstream and downstream regions surrounding each TE+ enhancer. TE classes are grouped as RNA-derived elements, including LTR, LINE, and SINE elements, and DNA-derived elements, including TIR elements. Asterisks indicate significance from chi-square goodness-of-fit tests comparing observed TE class composition (ns = not significant, *P < 0.01, **P < 0.001, ***P < 10^-6^). **(E)** Log_2_ enrichment of TE classes among TE+ enhancers relative to their matched local neighborhoods. Values show log_2_(observed/neighborhood) TE insert frequency for shadow and single enhancers, shown separately by tissue and for the pooled dataset. Filled circles indicate shadow enhancers, open circles indicate single enhancers, and diamonds indicate pooled values. Asterisks indicate FDR-adjusted significance for per-class observed-versus-neighborhood enrichment tests (*P < 0.01, **P < 0.001, ***P < 10^-6^). **(F)** (left) Example of a TE+ shadow enhancer set at the *Rbfox3* locus. Three shadow enhancers are TE+ from the same TE family, LTR/ERVL-MaLR, consistent with repeated co-option from a relatively homogeneous TE source. Right, RepeatMasker percent divergence from the consensus sequence is shown for the TE insertions overlapping the *Rbfox3* shadow enhancers. Similar divergence values provide a relative proxy for insertion age and are consistent with accumulation of these related TE insertions over a relatively narrow evolutionary interval. **(G)** Schematic illustrating candidate TE-mediated enhancer splitting events, in which TE sequence occupies most of the intervening space between two shadow enhancers, versus apparent non-splitting events, in which the TE appears to lie between two already separate enhancers. **(H)** Fraction of mouse shadow enhancer sets containing at least one candidate TE-mediated splitting event. **(I)** Example of a putative TE-mediated enhancer splitting event at the *Klf3* locus. A LINE element from the L1Md family lies between enhancer 1 and enhancer 2 within an eight-enhancer shadow enhancer set, consistent with a possible TE-mediated splitting event.

In the analysis of TE-derived fly enhancers, we found differences in the TE-class composition between shadow and single enhancers, which was partially explained by differences in the local genome neighborhoods. In contrast, mouse TE+ shadow and single enhancers showed a shared TE-class bias. Both groups were primarily derived from LTR and LINE elements, despite being located in SINE-rich local neighborhoods (Figure 6D). This pattern was consistent across forebrain, heart, and limb enhancers, even though these tissues use distinct transcription factor inputs, and was also recovered using an alternative measure of TE composition (Figure 6E; Supplementary Figure 15). This suggests mouse TE co-option is shaped by TE class but is not specific to shadow enhancer architectures. For example, three of 23 enhancers in the *Rbfox3* forebrain shadow set are TE-derived, all from the LTR/ERVL-MaLR family, with similar RepeatMasker divergence values consistent with repeated co-option from a common TE-family pool (Figure 6F).

Together, these results suggest that TE co-option contributes to mouse shadow enhancer sets, but is not a shadow-specific route of enhancer birth. Instead, TE-derived enhancers occur at similar frequencies in shadow and single enhancer architectures, with both groups showing preferential contribution from LTR and LINE elements rather than simply reflecting the local TE neighborhood.

### TE-mediated splitting is rare in mouse shadow enhancer sets

Finally, we tested whether TEs may contribute to mouse shadow enhancer birth through TE splitting (Figure 6G). Using the same criteria as in fly, we identified 162 candidate TE splitting hits across 9,051 shadow enhancer sets, corresponding to only 1.8% of mouse shadow sets (Figure 6H). One candidate occurs near *Klf3*, where an L1Md LINE lies between two enhancers in an eight-enhancer shadow set (Figure 6I). Together with the fly results, this low frequency suggests that TE-mediated splitting is uncommon in both genomes, whereas TE co-option and duplication contribute more frequently to shadow enhancer architectures.

## Discussion

Shadow enhancers are widespread features of developmental regulatory landscapes. There are many potential routes of shadow enhancer birth, which can shape the regulatory logic and function of these common DNA elements. Here, we tested whether shadow enhancer sets in *Drosophila melanogaster* and mouse show sequence signatures consistent with three possible birth mechanisms: duplication of existing enhancers, TE co-option, and TE-mediated splitting of ancestral regulatory elements. Overall, our results suggest that shadow enhancer birth is not dominated by a single one of these evolutionary mechanisms. Shadow enhancers appear to originate using a mixture of mechanisms, with duplication accounting for a substantial subset of cases, TE co-option contributing to both shadow and non-shadow enhancer birth, and TE-mediated splitting appearing rare genome-wide.

### Genome architecture shapes the shadow birth routes in fly and mouse

By applying a common analytical framework to fly and mouse developmental shadow enhancer sets, we found shared and species-specific patterns. Duplication-based shadow sets were more common in the mouse genome than the fly genome. TE+ enhancers were also more common in mouse shadow sets than fly shadow sets, while TE-mediated splitting candidates were rare in both species. The stronger duplication- and TE-associated signals in the mouse genome likely reflect differences in overall genome architecture. The mouse genome is roughly ten times larger than the *Drosophila* genome^34,35^ and contains more of the sequence substrates relevant to our birth models. About 1.4% of the *D. melanogaster* genome arose from relatively recent segmental duplications, compared to 4.94% of the mouse^29,36^. Similarly, TE content is ∼21.7–22.8% in the *D. melanogaster* genome, compared with 37.5% in the mouse genome^32,34^. These differences may increase the opportunity for the duplication of noncoding regulatory sequences and TE co-option in the mouse genome and support a model in which the relative contribution of shadow enhancer origin routes depends in part on genome architecture.

### TE co-option contributes enhancer material, but is not shadow-specific

Our data do not support the idea that TE co-option is specific to shadow enhancer birth. In both genomes, TE+ enhancer frequency is not significantly different between shadow and single enhancers (Figure 3C; Figure 6C). This finding is consistent with a previous analysis of human and mouse shadow enhancers^37^. However, we found a lower fraction of TE+ enhancers (7.5%) than this previous study, in which 37% of mouse enhancers were called TE+. This difference is likely due to a different method for detecting shadow sets based on enhancer RNA correlation with gene expression; analysis of enhancers active in different tissues; and a ∼10% overlap threshold to call an enhancer TE+ versus our 20% threshold. Other work to identify TE-derived human and mouse enhancers found that the fraction of TE+ enhancers depends on the threshold used and by the cell- or tissue-type considered^38^. Together, both our work and that of Barth et al. suggest that TEs contribute to enhancer birth generally, but do not preferentially generate redundant enhancer architectures.

### TE co-option class bias is more apparent in mouse, whereas fly TE co-option reflects the local genomic neighborhood

The TE co-option data in fly and mouse suggest that TE class identity and local genomic context can both shape co-option. In the mouse genome, both shadow and single TE+ enhancers were preferentially derived from LTR and LINE elements even though their surrounding neighborhoods were SINE-rich, suggesting that co-option in the mouse genome is not simply proportional to local TE availability (Figure 6D,E). This pattern is consistent with prior work showing that retrotransposons contribute to tissue-specific and rapidly evolving regulatory regions in mammals^39–42^. In the fly genome, by contrast, TE-class composition differed between shadow and single enhancers, and these differences were partly explained by local neighborhood composition (Figure 3D,E). We further hypothesized that the remaining differences in TE-class composition were due to the TFBS content in TE sequences, but found little evidence to support this idea (Supplementary Figure 9). These results and other studies paint a complex picture, in which the likelihood of TE co-option is modulated by several factors, including local TE availability, TE sequence content, developmental chromatin environment, and subsequent evolutionary refinement^43–45^.

### Birth route may influence, but does not determine, enhancer regulatory logic

Shadow enhancers are defined by their ability to drive identical or overlapping expression patterns. Some birth mechanisms, e.g. duplication, immediately give rise to highly similar or identical enhancers, while others, e.g. de novo evolution, require convergent evolution to achieve redundancy. Further, redundant activity does not imply identical regulatory grammar. Some shadow enhancer pairs rely on distinct TF inputs, which can allow for less noisy gene expression from a shadow set^8,12,46^.

Enhancer duplication provides the clearest case in which the birth mechanism predicts regulatory architecture, and our duplication-hit pairs show elevated TFBS similarity relative to non-hit shadow pairs. However, duplicated pairs also span a broad range of TFBS similarity at comparable sequence identities, suggesting multiple post-duplication trajectories ranging from strict conservation to regulatory rewiring (Figure 2C). This is consistent with models in which enhancer function can be maintained despite extensive TFBS turnover^26,27,47,48^.

TE co-option provides a complementary test of whether birth route predicts regulatory logic. If TE regulatory cargo predicted enhancer co-option, enriched TE classes have similar motifs to TE+ enhancers. Instead, fly TE co-option by class showed weak correlation with motif prevalence, while mouse TE co-option showed similar class biases across three tissues with distinct TF inputs (Figures 3D and 6E). Consistent with Barth et al., where most TE-derived mouse redundant enhancer pairs came from distinct TE families, these results suggest that TE cargo alone does not determine which TEs become enhancers^37^.

### Limitations and implications

Using an enhancer-level tally, the three mechanisms examined here account for only a minority of shadow enhancers: ∼23% in fly and ∼37% in mouse. Some shadow enhancers likely arose through distinct routes, including de novo TFBS gain or co-option of existing enhancers to new target genes, but ancient duplications, older TE exaptations, or ancestral regulatory relationships may no longer retain recognizable sequence signatures. However, given that the elements have turned over sufficiently to obscure their origins, we can classify them as de novo-like: functionally overlapping elements whose birth signatures are no longer detectable. This interpretation fits the broader view that enhancers can emerge from diverse sequence substrates and that similar regulatory outputs can evolve through different combinations of binding sites^3,13,22–24,27,49–51^. Together, our results add to a growing body of work emphasizing the flexibility of enhancer sequence-function relationships and suggest that overlapping regulatory activity can arise through convergent or stepwise assembly of enhancer logic, rather than solely through inheritance of a common regulatory template.

## Methods

### Shadow enhancer datasets and genome assemblies

*D. melanogaster* shadow enhancer annotations were taken from two published datasets: early developmental enhancers assayed in^25^ and mesoderm-associated enhancers defined from transcription factor binding and gene expression data^4^. Mouse shadow enhancer annotations were taken from the Osterwalder et al. developmental enhancer dataset^5^. In each species, shadow sets were defined as gene–tissue groups, or gene–tissue/time groups for mouse, containing at least two enhancers assigned to the same gene and activity context. Redundant entries with identical enhancer membership were collapsed so that each retained entry represented a unique shadow set. The final datasets contained 420 unique fly shadow enhancer sets comprising 1,122 unique enhancers and 9,051 unique mouse shadow enhancer sets comprising 19,166 unique enhancers. Single enhancers were retained from the same source datasets for TE co-option comparisons and included 6,673 fly single enhancers and 6,363 mouse single enhancers. Analyses were performed using the dm6 assembly for fly and the mm10 assembly for mouse.

### Shadow enhancer-body homology analysis for duplication screen

BLAST^52^ searches were performed using the following parameters: `-evalue 1 -word_size 10 -gapopen 5 -gapextend 2 -reward 2 -penalty -3 -dust yes`. Because fly and mouse genomes differ in size, repeat content, and background sequence similarity, BLAST significance thresholds were calibrated separately for each species. For each genome, a null distribution of BLAST E-values was generated from pairwise comparisons of all shadow enhancers. Species-specific E-value 90% cutoffs were then selected from these null distributions and applied to the corresponding shadow enhancer dataset. The resulting enhancer-body cutoffs were E ≤ 10^-4^ for fly and E ≤ 10^-8^ for mouse. Enhancer pairs with at least one significant enhancer-body BLAST alignment were classified as enhancer-hit pairs.

### TFBS coverage of fly BLAST hits

BLAST-hit sequences were scanned for predicted TFBS. All available *D. melanogaster* transcription factor motifs were downloaded from JASPAR (n = 162 motifs) ^53^, and FIMO^54^ was used to scan BLAST-hit sequences using a significance threshold of P < 10^-3.^. For each BLAST hit, TFBS coverage was calculated as the fraction of aligned base pairs overlapped by one or more FIMO motif calls.

### Duplication detection horizon

To estimate the detection horizon of the BLAST-based duplication screen, we used the stringent set of 370 “deeply conserved” *D. melanogaster* S2-cell enhancers from Arnold et al.^22^, which were active across the original five-species *Drosophila* comparison: *D. melanogaster*, *D. yakuba*, *D. ananassae*, *D. pseudoobscura*, and *D. willistoni*. Orthologous enhancer sequences were extracted from UCSC Multiz insect whole-genome alignments using the D. melanogaster enhancer coordinates^22,55^. We first applied the same BLAST parameters used in the fly shadow enhancer duplication screen to compare homologous functional enhancers across the Arnold et al. five-species panel. We then extended the analysis to additional aligned species, including *D. sechellia*, *D. simulans*, *D. erecta*, *D. persimilis*, *D. mojavensis*, *D. virilis*, *D. grimshawi, Musca domestica*, *Anopheles gambiae*, and *Tribolium castaneum*. For each species, we calculated the fraction of orthologous enhancer sequences with at least one significant BLAST hit to the corresponding *D. melanogaste*r enhancer.

### Flanking-sequence homology screen and duplication cohort definitions

For each enhancer pair within a shadow set, 2 kb 5′ and 3′ flanking intervals were extracted from the reference genome. Flanks were truncated at chromosome boundaries and were not allowed to include enhancer bodies from the same shadow set. For enhancer pairs on the same chromosome separated by less than 4 kb, inner flanks were adjusted to the intervening sequence between the two enhancers, retaining only sequence contiguous with each enhancer edge. If no intervening sequence remained, the corresponding inner flank was omitted.

We generated a species-specific null distribution of flank-alignment E-values from randomly paired flanks and used the null-calibrated 90% E-value cutoff as the flank-alignment threshold. Each observed pair was tested across four flank orientations: left-left (LL), right-right (RR), left-right (LR), and right-left (RL). Single-flank support required one significant flank hit. Paired-flank support required two orientation-consistent hits, either LL+RR or LR+RL. Direct inner-flank comparisons were excluded for adjacent same-chromosome pairs when they would compare overlapping or shared intervening sequences. Enhancer-body hit pairs were annotated as having no, single-flank, or paired-flank support. Pairs without enhancer-body hits but with orientation-consistent paired-flank support were classified as flank-only pairs.

### Duplication proportion by shadow set size

Shadow sets were grouped into 2-enhancer, 3-enhancer, and ≥4-enhancer bins. For each bin, we calculated the fraction of enhancer memberships participating in at least one duplication-hit pair. Significance was assessed against 100 size-matched randomized datasets generated by sampling enhancers together with their associated flanking sequences while preserving the observed shadow set size distribution. Randomized sets were analyzed using the same enhancer-body and flank-comparison pipeline, including the same BLAST parameters and species-specific E-value cutoffs. Observed duplication-hit proportions were compared with size-matched randomized null distributions by calculating a Z-score from the mean and standard deviation of the randomized replicates, with two-sided P-values obtained from the normal approximation. Pairwise differences in duplication-hit proportions between set-size bins were assessed using Bonferroni-corrected two-proportion Z-tests.

### TFBS analysis of duplication-hit shadow enhancer pairs

The TFBS similarity between enhancer pairs was calculated using 14 mesoderm TF motifs from JASPAR database^53^: Biniou (Bin), Twist (Twi), Mothers against dpp (Mad), Pangolin (Pan), Pannier (Pnr), Scalloped (Sd), Tinman (Tin), Snail (Sna), Dorsal (Dl), Even-skipped (Eve), Ladybird early (Lbe), Tailup/islet (Tup), Bagpipe(Bap), and Ladybird late (Lbl). No-hit pairs were defined as shadow enhancer pairs lacking both enhancer-body BLAST support and paired-flank support. Pairwise TFBS similarity was quantified using the Jaccard metric, which accounts for both motif identity and motif count. Statistical comparisons between duplication-hit and no-hit pairs were performed using Mann-Whitney U test.

Global sequence identity was calculated from pairwise Needleman–Wunsch alignments of full enhancer sequences and reported as percent identity. The relationship between sequence identity and TFBS similarity was modeled by ordinary least-squares linear regression. Residuals from this model were standardized as Z-scores to identify enhancer pairs with higher- or lower-than-expected TFBS similarity given their overall sequence identity.

### TE annotation

TE annotations were generated or obtained for each genome. For *D. melanogaster,* TEs were annotated in dm6 using RepeatMasker v4.1.5^56^ with the *Drosophila* RepBase library^57,58^. To improve detection of unannotated or divergent repeats, de novo repeat discovery was also performed using RepeatModeler v2.0.3 ^59^, and the resulting custom library was used for a second RepeatMasker pass. Fragmented TE hits were merged into full-length elements using OneCodeToFindThemAll^60^. Redundant TE calls from RepeatMasker and RepeatModeler were merged when elements of the same TE type overlapped by at least 80%, using the outermost 5′ and 3′ coordinates to define the final insertion. The resulting fly TE annotation covered 20.8% of the dm6 assembly. For mouse, TE annotations were obtained from the UCSC mm10 RepeatMasker annotation^55,56^.

### Identification of TE+ enhancers

Enhancers were classified as TE-derived, or TE+, if at least 20% of the enhancer length overlapped annotated TE sequence. Enhancers that did not meet this threshold were classified as TE−. For enhancers overlapping more than one TE, all overlapping TE annotations were retained for class-level analyses.

### Fixation of enhancer-associated TEs in *Drosophila*

Following the TE fixation analysis of Rech et al., we evaluated our enhancer-associated TE insertions across the ten long-read *D. melanogaster* strain assemblies from that study^32^. For each ISO1 TE insertion overlapping at least 20% of an enhancer, we extracted the TE sequence plus 500 bp of flanking sequence on each side and mapped these sequences to each strain assembly using minimap2 v2.9 with the splice preset^33,61^. Flanking alignments were used as anchors to infer whether the TE insertion was present at the orthologous locus. Calls were retained only when both flanks mapped unambiguously. Enhancer-associated TEs were classified as fixed when present in at least 90% of strains, and not fixed otherwise. Cases in which orthology could not be confidently assessed in at least eight lines were classified as null.

### TE-class and local-neighborhood analyses

To determine whether TE-derived enhancers reflect local TE availability or preferential co-option of specific TE families, we compared TE-class distributions among TE+ shadow enhancers, TE+ single enhancers, local genomic neighborhoods, and the genome-wide TE background. For each TE+ enhancer, local neighborhood was defined as the enhancer interval plus 50 kb upstream and 50 kb downstream. TE families were collapsed into four broad classes: LTR, LINE, SINE for RNA group and Helitron/RC and TIR for DNA group. The mouse genome is not known to have Helitron/RC class and so that class was excluded for mouse genome. For each enhancer class, observed TE-class composition was calculated from unique enhancer–TE overlap pairs, and the local background was calculated from TE insertions overlapping the corresponding enhancer-centered neighborhood windows. TE-class composition differences were assessed using the chi-square goodness-of-fit test. TE-class enrichment was calculated as:

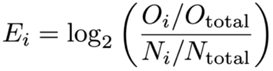

where O_i_ is the number of observed enhancer–TE overlap pairs from TE class i, O_total_ is the total number of enhancer–TE overlap pairs, N_i_ is the number of local-neighborhood TE insertions from TE class i, and N_total_ is the total number of local-neighborhood TE insertions. Statistical significance for each TE class was assessed using a two-sided one-sample proportion Z-test comparing the observed fraction of enhancers overlapping TE class i to the corresponding local-neighborhood TE-class proportion. P-values were adjusted across TE-class tests with Benjamini–Hochberg procedure.

### TE TFBS analysis

To test whether TE co-option patterns reflected TE class-specific regulatory cargo, we scanned shadow enhancers, single enhancers, and TE sequences for the same Jaspar database 14 mesoderm-associated TF motifs used in the enhancer TFBS similarity analysis using FIMO, 10^-3^ cutoff: Biniou (Bin), Twist (Twi), Mothers against dpp (Mad), Pangolin (Pan), Pannier (Pnr), Scalloped (Sd), Tinman (Tin), Snail (Sna), Dorsal (Dl), Even-skipped (Eve), Ladybird early (Lbe), Tailup/islet (Tup), Bagpipe (Bap), and Ladybird late (Lbl). For each enhancer group or TE class, we calculated the fraction of sequences containing at least one FIMO-predicted match to each motif. Similarity between enhancer and TE motif profiles was quantified using Pearson correlation across the 14 motifs.

### TE splitting analysis

For each shadow enhancer set, inter-enhancer sequences were defined as the genomic intervals between adjacent enhancers. These “spacer” sequences were intersected with the species-specific TE annotation to calculate TE coverage of the spacer and remaining non-TE sequence. Candidate splitting events were defined using two criteria: either TE sequence occupied at least 80% of the spacer, or less than 500 bp of non-TE sequence remained within a TE-containing spacer.

## Acknowledgements

This work was supported by NIH award R01HD095246 (to ZW) and a Kilachand Multicellular Design Program Fellowship and a BU Nano Cross-Disciplinary Fellowship (to JN). The content is solely the responsibility of the authors and does not necessarily represent the official views of the National Institutes of Health. The authors wish to thank Sima Tahmouresie and Aman Burji for initial work on this project and Eric Gerber for comments on the manuscript.

