## Supplementary Figures and Tables for "Genome architecture shapes the evolutionary origins of redundant enhancers in fly and mouse"

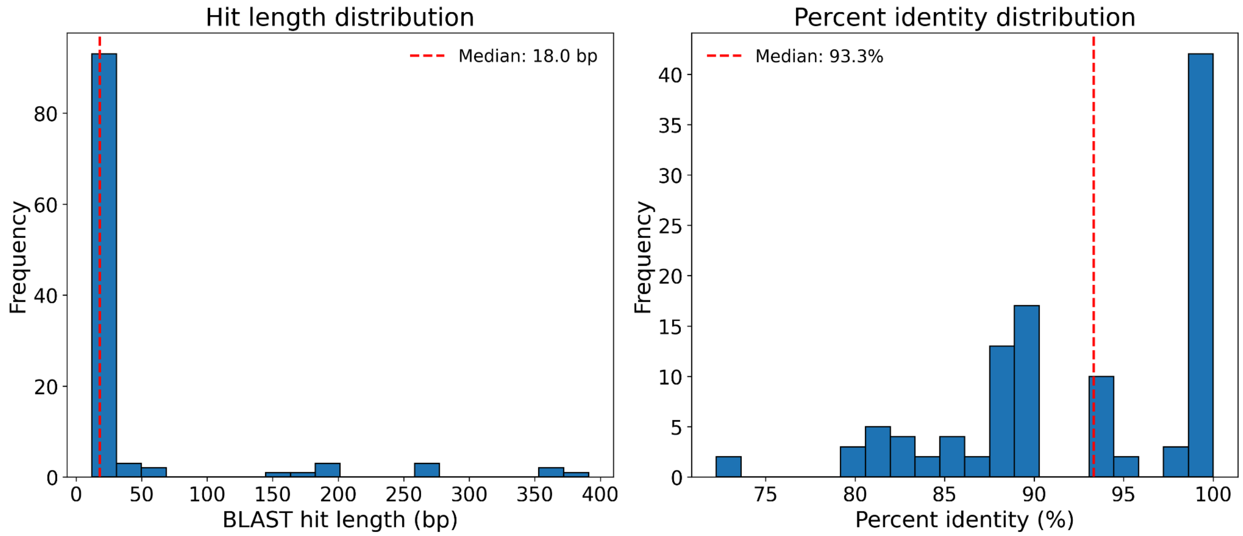


**Supplementary Figure 1. Length and percent identity of fly enhancer BLAST hits**. Distributions of BLAST hit length and percent identity are shown for fly enhancer-body hits. Dashed lines indicate medians.

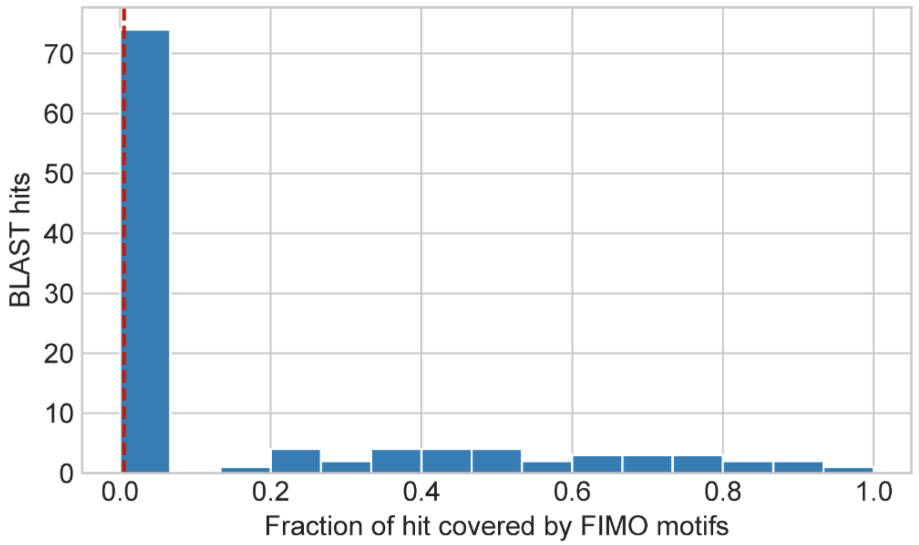


**Supplementary Figure 2. Transcription factor binding site coverage of fly BLAST hits.** Distribution of TFBS motif coverage across fly enhancer-body BLAST-hit sequences. Median motif coverage is 0.0%.

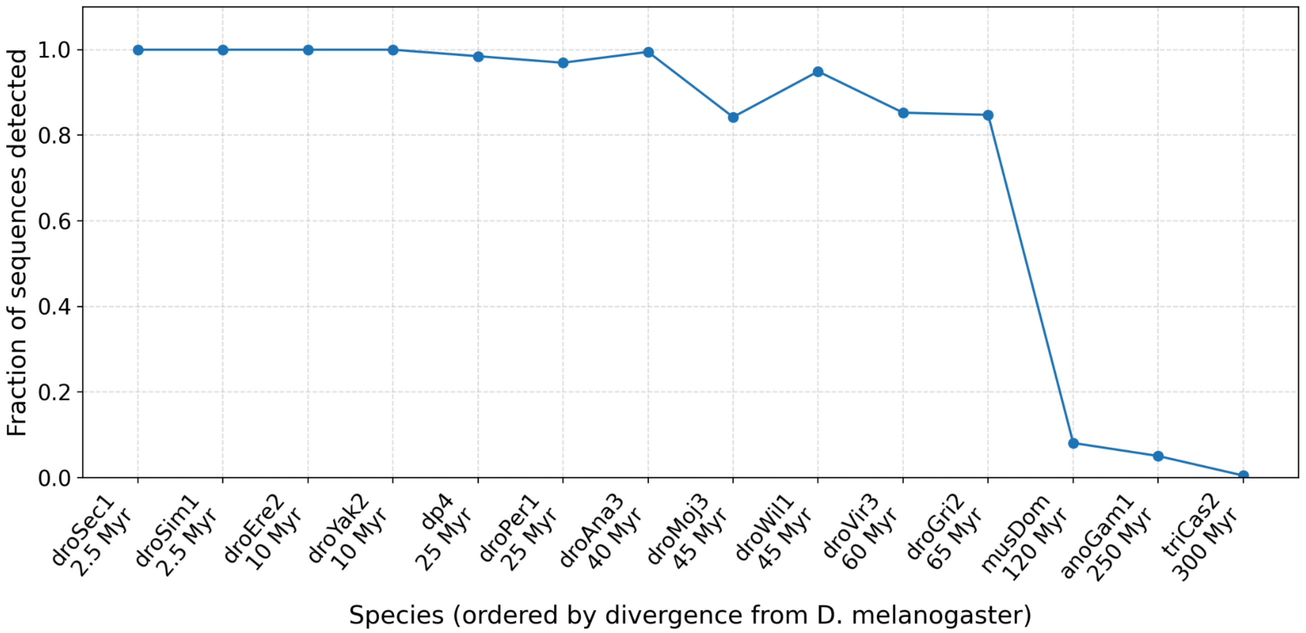


**Supplementary Figure 3. Sensitivity of the BLAST-based enhancer-homology screen across evolutionary distances.** To estimate the detection horizon of the BLAST approach, the same BLAST parameters used in the fly shadow enhancer duplication screen were applied to functionally validated homologous enhancers from Arnold et al.[21]. Orthologous enhancer sequences were extracted from UCSC Multiz insect whole-genome alignments using the *D. melanogaster* enhancer coordinates[22,55]. The analysis first used the original Arnold et al. five-species panel (*D. melanogaster*, *D. yakuba*, *D. ananassae*, *D. pseudoobscura*, and *D. willistoni*; n = 370 enhancers), then extended to additional aligned species. For each species, we calculated the fraction of orthologous enhancer sequences with at least one significant BLAST hit to the corresponding *D. melanogaster* enhancer. The screen recovered most homologous enhancer sequences across recent-to-intermediate divergence times, but recovery at deep divergence fell to ~5%, matching the empirically calibrated false-positive rate. Thus, at deep divergence, BLAST recovery is not above background, indicating that the duplication screen is most sensitive to recent-to-intermediate homology. Species abbreviations follow the plotted assembly labels: droSec1, *D. sechellia*; droSim1, *D. simulans*; droEre2, *D. erecta*; droYak2, *D. yakuba*; dp4, *D. pseudoobscura*; droPer1, *D. persimilis*; droAna3, *D. ananassae*; droMoj3, *D. mojavensis*; droWil1, *D. willistoni*; droVir3, *D. virilis*; droGri2, *D. grimshawi*; musDom, *Musca domestica*; anoGam1, *Anopheles gambiae*; and triCas2, *Tribolium castaneum*.

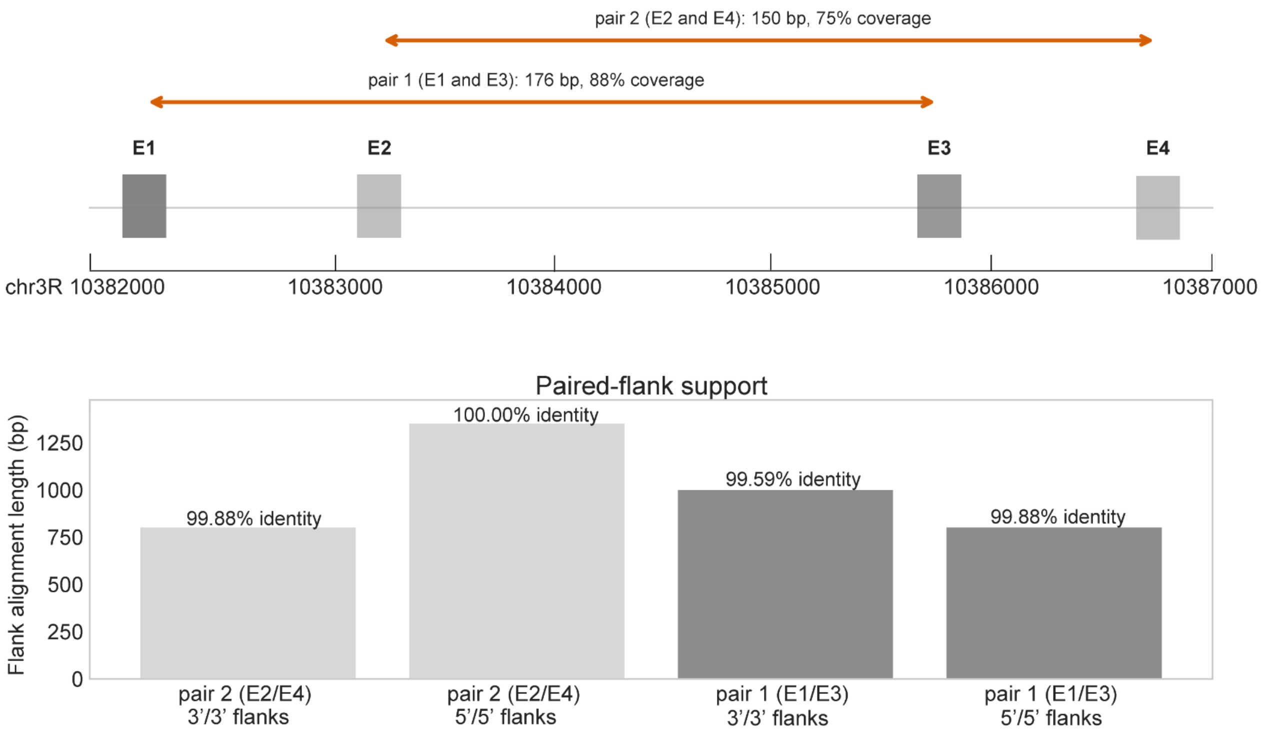


**Supplementary Figure 4. The CG6345 shadow enhancers have strong evidence of a tandem duplication event.** Top, four CG6345 shadow enhancers are shown in locus order, with duplicated enhancer pairs connected by arrows. Pair 1 (E1 and E3) is shown in dark grey. Pair 2 (E2 and E4) is shown in light grey. Bottom, paired-flank alignments summarize the support for each duplicated enhancer pair.

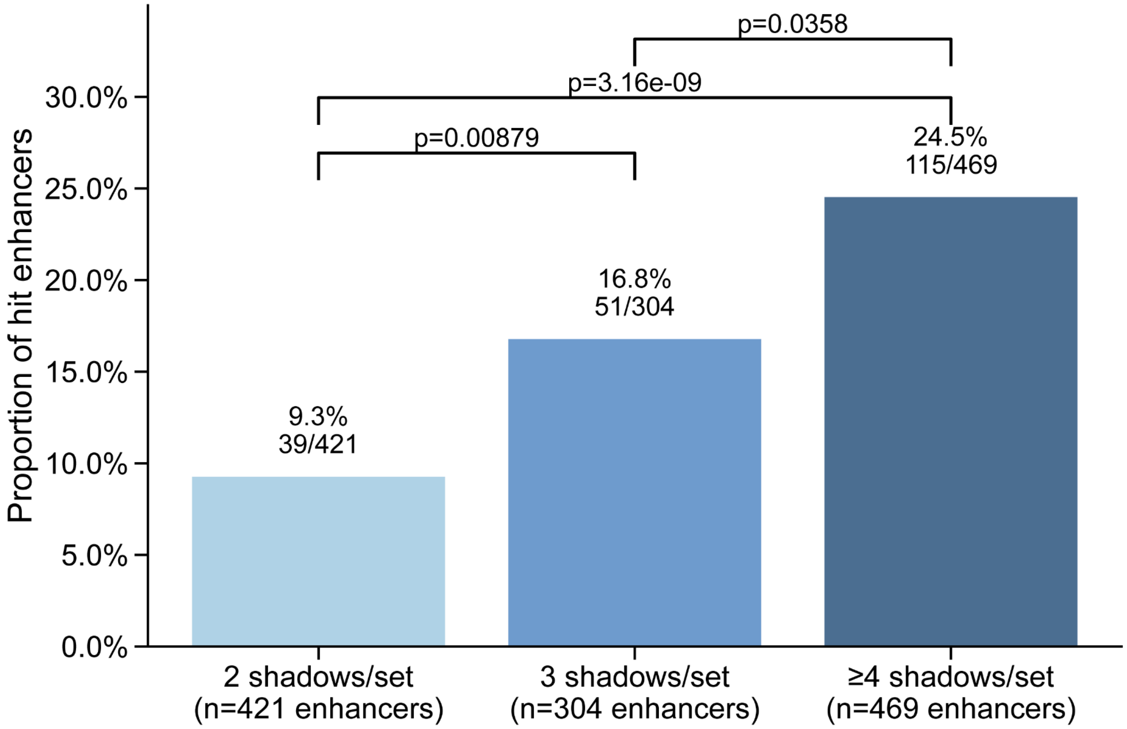


**Supplementary Figure 5. Fly duplication hit enhancer percentage by shadow set size bins.** The observed fraction of fly enhancers participating in at least one duplication-hit pair is shown for shadow sets containing 2, 3, or ≥4 enhancers. Pairwise two-sided Z-tests performed with Bonferroni correction.

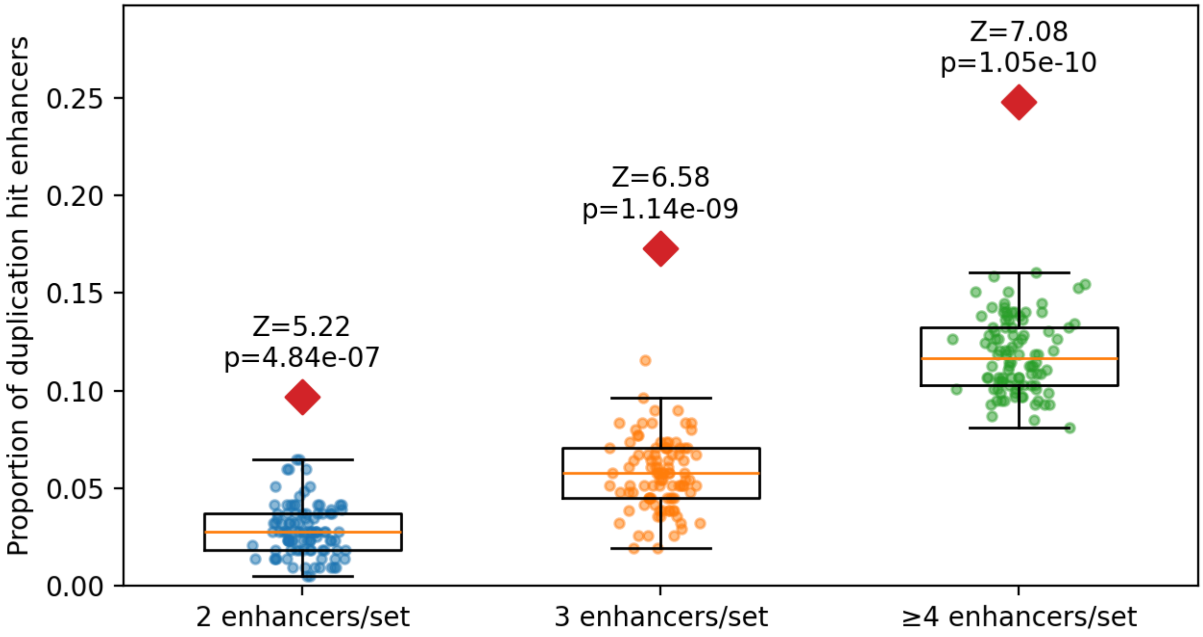


**Supplementary Figure 6. Size-matched randomized control for fly duplication enrichment by shadow-set size.** Observed fractions of duplicated enhancers in fly shadow sets are shown in red diamonds for each shadow-set size bin: 2 enhancers/set, 3 enhancers/set, and ≥4 enhancers/set. Distribution of 100 size-matched randomized control replicates is shown with the box-and-whisker plots. Z-scores and p-values were calculated by comparing the observed fraction with the corresponding randomized-control distribution for each size bin.

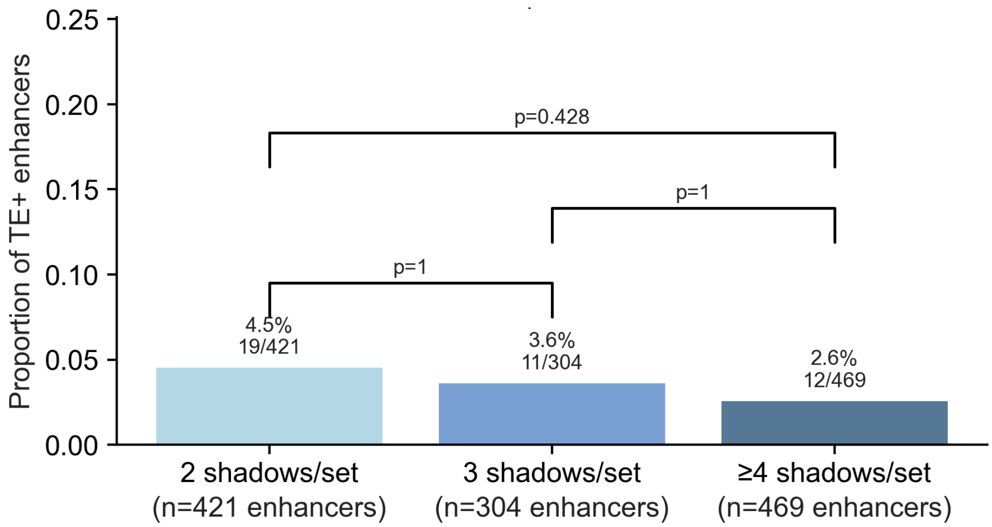


**Supplementary Figure 7. Fly TE+ enhancers by shadow set size.** Bars show the fraction of enhancers that are TE+ in shadow sets containing 2, 3, or ≥4 enhancers. Pairwise two-sided Fisher exact tests performed with Bonferroni correction.

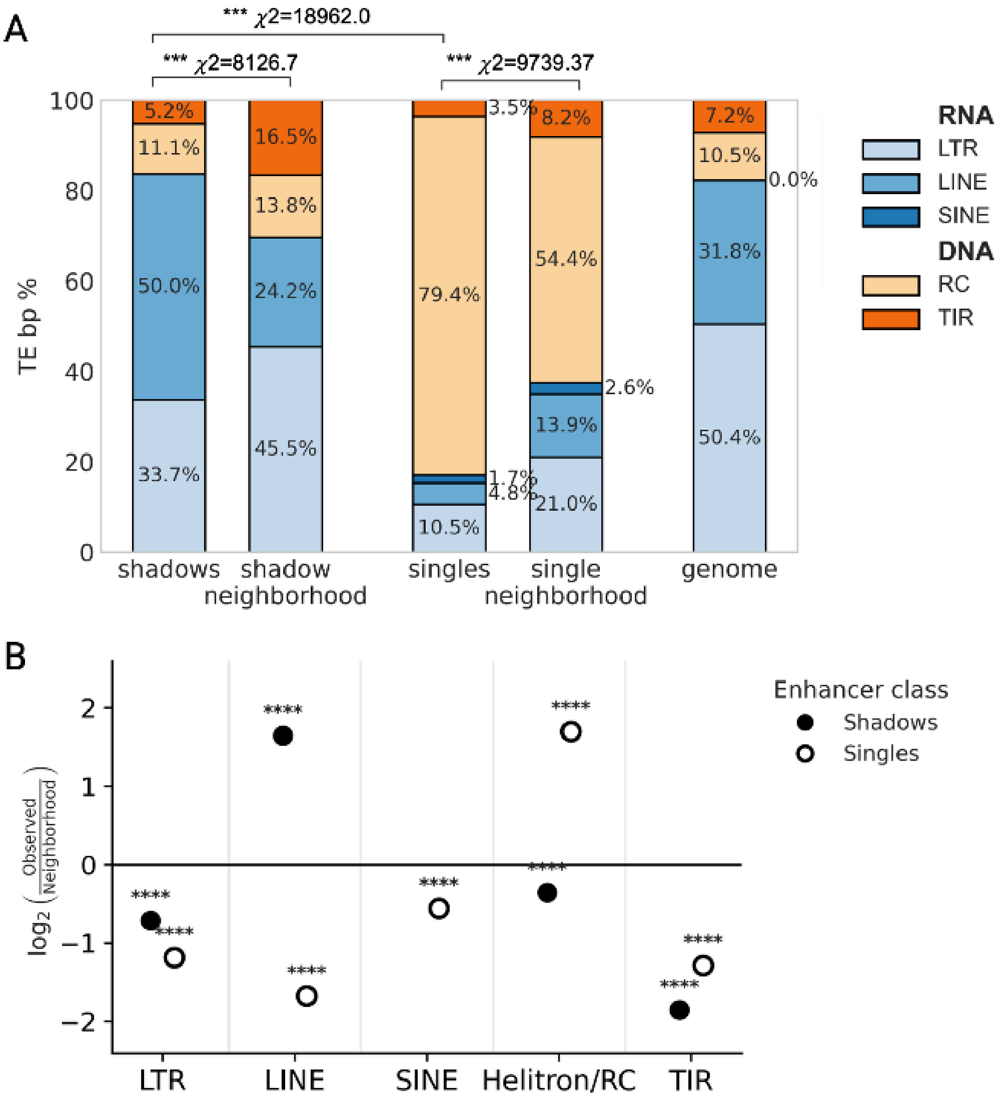


**Supplementary Figure 8. Fly TE class composition among TE co-opted shadow and single enhancers, analyzed by basepair composition. (A)** Stacked bars show the base-pair composition of overlapping TE+ shadow enhancers, TE+ single enhancers, their matched local neighborhoods, and the genome. This analysis is similar to Figure 3D and E, but analyzed by number of basepairs, instead of numbers of insertion events. TE classes are grouped as RNA-derived elements, including LTR, LINE, and SINE elements, and DNA-derived elements, including Helitron/RC and TIR elements. Asterisks indicate significance from chi-square goodness-of-fit tests comparing observed TE class composition to the corresponding local neighborhood or genome background. **(B)** $Log_2$ observed-over-neighborhood enrichment of TE classes among TE+ shadow and single enhancers. Values show $log_2$(observed/neighborhood) TE base-pair frequency for shadow and single enhancers. Filled circles indicate shadow enhancers and open circles indicate single enhancers. Asterisks indicate FDR-adjusted significance for per-class observed-versus-neighborhood enrichment tests.

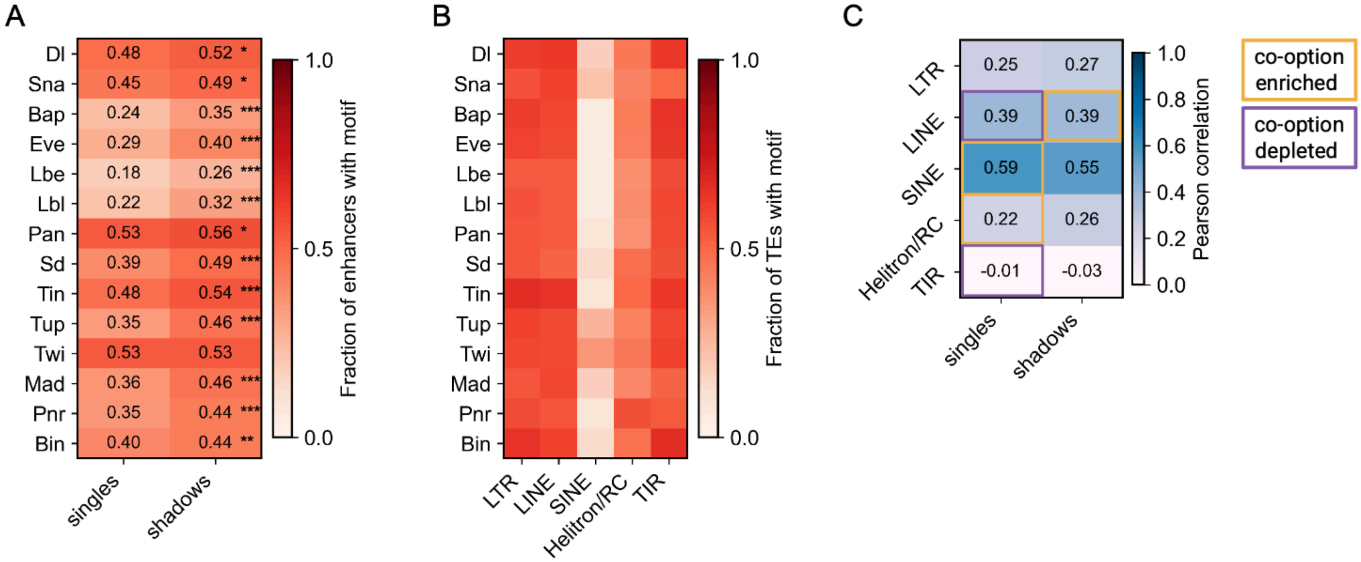


**Supplementary Figure 9. Motif profiles of TE classes do not fully explain co-option biases in shadow and single TE+ enhancers. (A)** TFBS prevalence heat map showing the fraction of shadow and single enhancers containing at least one predicted motif for each of 14 mesoderm-associated transcription factors. Asterisks indicate FDR-adjusted significance from per-TF z-score proportion tests. (ns = not significant, *P < 0.01, **P < 0.001, ***P < $10^{-6}$). **(B)** TFBS prevalence heat map showing the fraction of TE sequences within each TE class containing at least one predicted motif for each of the same 14 transcription factors.(**C**) Correlation of shadow and single enhancers with TE class prevalence maps. Yellow boxes indicate when a specific TE class was enriched compared to background in co-option and purple represents a TE class that was depleted compared to background (Figure 3E). TE-class motif-prevalence profiles showed only modest correlations with shadow and single enhancer motif profiles, and these correlations did not consistently predict co-option enrichment. For example, DNA/RC motif prevalence profiles are weakly correlated with single enhancer profiles ($r$ = 0.22), but are actually slightly more correlated with shadow enhancer profiles ($r$ = 0.26). LINEs, which are more commonly co-opted for shadow enhancers than single enhancers, showed the same correlation ($r$ = 0.39) for both types of enhancers.

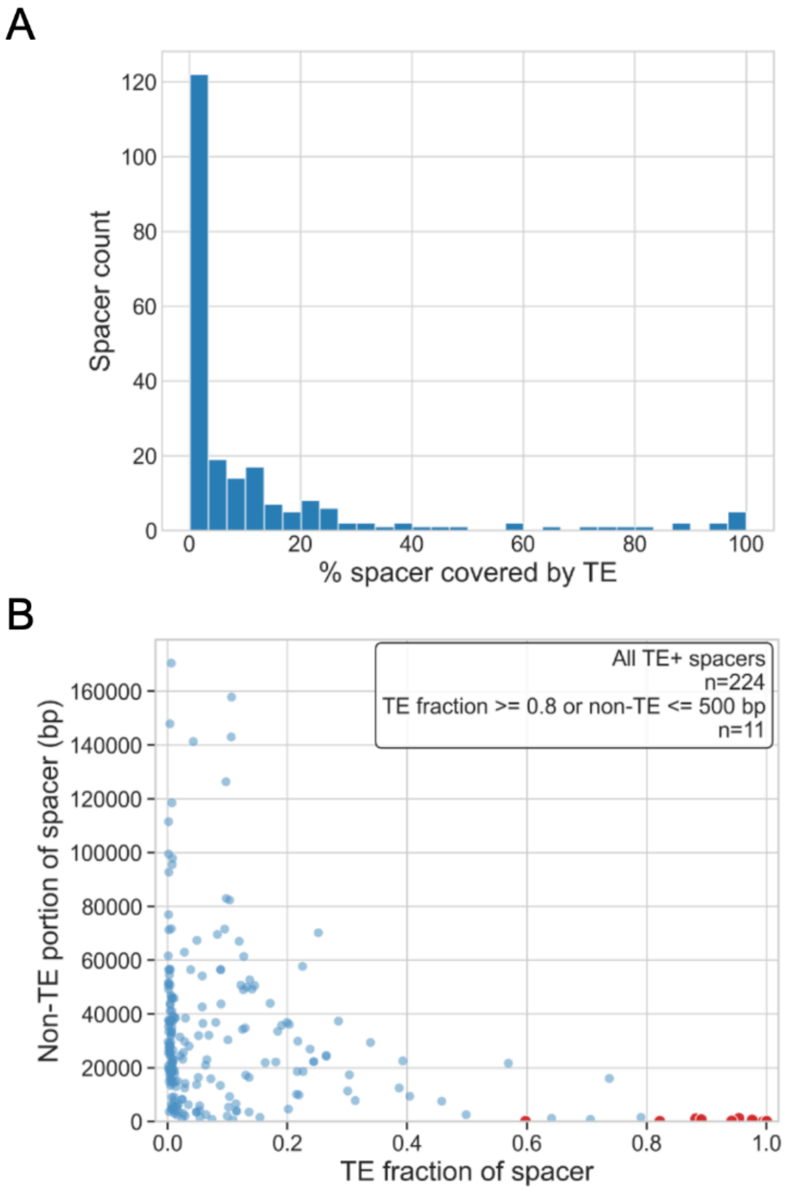


**Supplementary Figure 10. Fly candidate TE-splitting spacer sequences represent high-coverage TE-containing inter-enhancer regions.** To identify candidate TE-mediated enhancer splitting events, we examined the intervening sequence between enhancer pairs within each shadow enhancer set**.** Spacers were intersected with the TE annotation to calculate TE coverage of the spacer, and remaining non-TE sequence **(A)** Distribution of TE coverage of TE+ inter-enhancer spaces. (**B**) TE+ high-coverage spacers were defined as TE-containing spacer sequences in which TE sequence occupies ≥80% of the intervening region or in which <500 bp of non-TE sequence remains (red points).

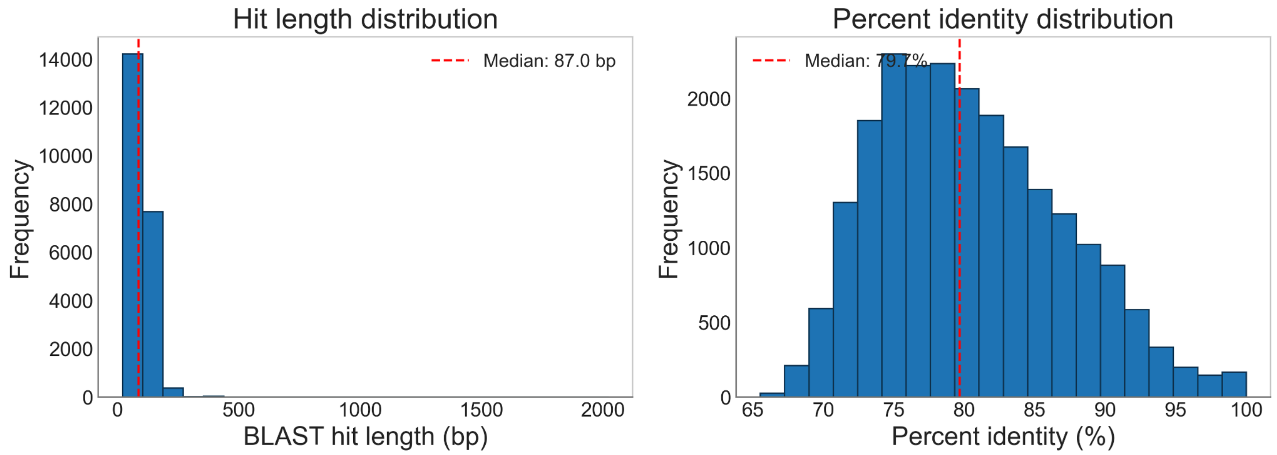


**Supplementary Figure 11. Mouse length and percent identity of enhancer BLAST hits**. Distributions of mouse BLAST hit length and percent identity are shown for mouse enhancer-body hits. Dashed lines indicate medians.

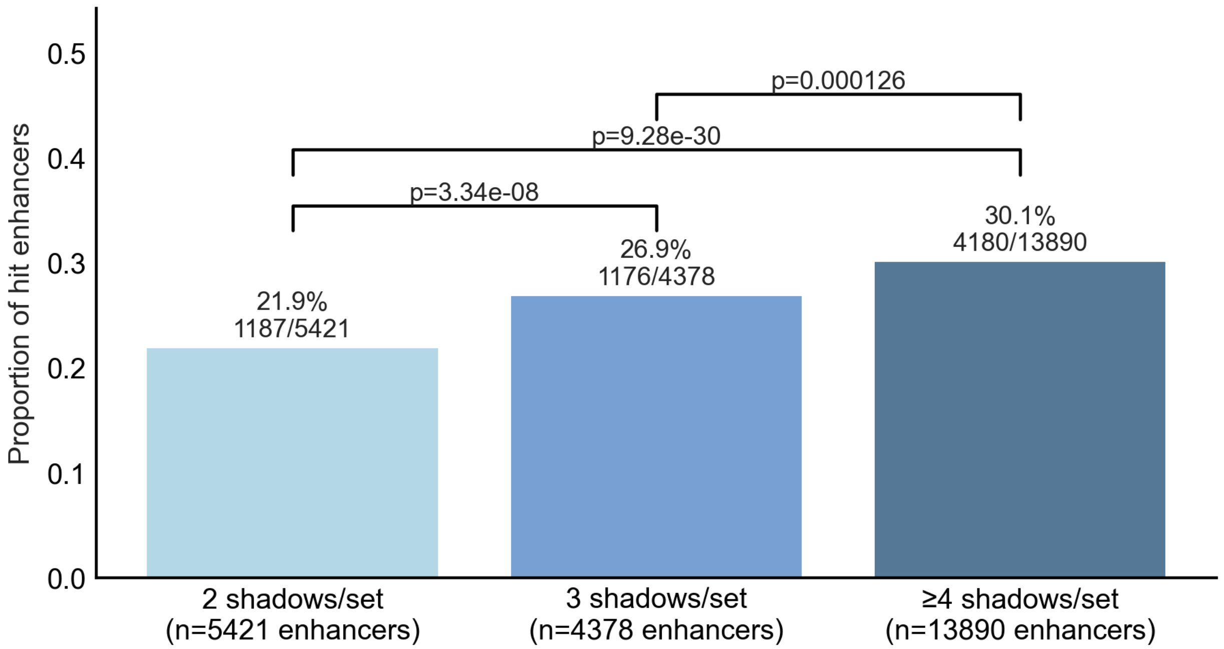


**Supplementary Figure 12. Mouse duplication hit enhancer percentage by shadow set size bin.** The observed fraction of mouse enhancers participating in at least one duplication-hit pair is shown for shadow sets containing 2, 3, or ≥4 enhancers. Pairwise two-sided Z-tests performed with Bonferroni correction.

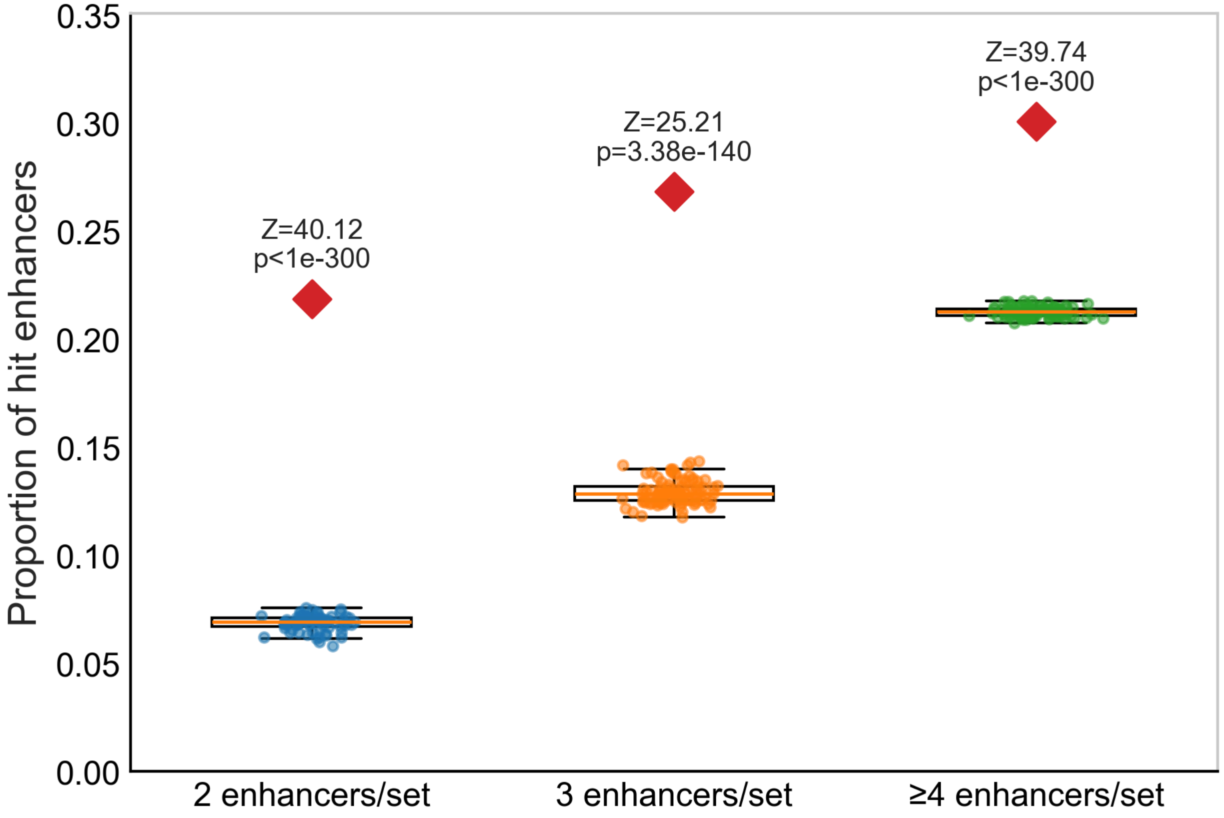


**Supplementary Figure 13. Mouse size-matched randomized control for duplication enrichment by shadow-set size.** Observed fractions of duplicated enhancers in mouse shadow sets are shown in red diamonds for each shadow-set size bin: 2 enhancers/set, 3 enhancers/set, and ≥4 enhancers/set. Distributions from 100 size-matched randomized control replicates as box-and-whisker plots. Z-scores and p-values were calculated by comparing the observed fraction with the corresponding randomized-control distribution for each size bin.

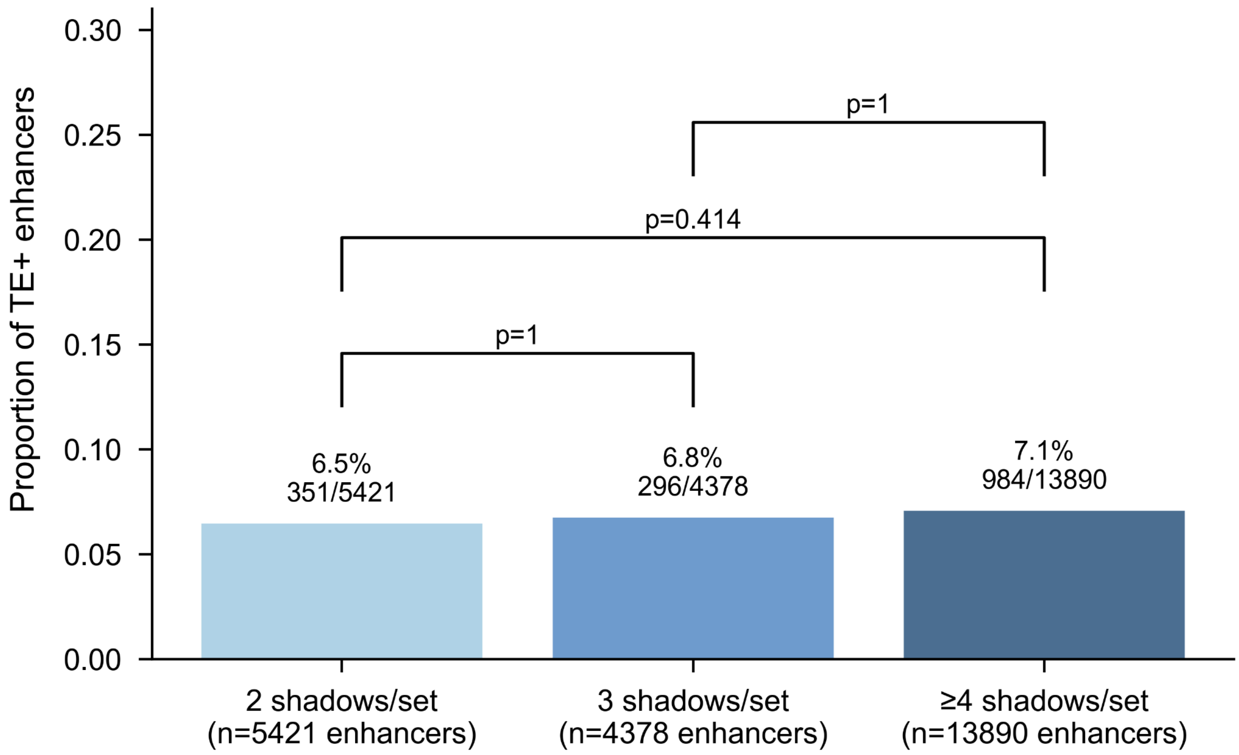


**Supplementary Figure 14. Mouse TE+ enhancers by shadow set size**. Bars show the fraction of enhancers that are TE+ in shadow sets containing 2, 3, or ≥4 enhancers. Brackets mark two-sided Z-tests with Bonferroni correction.

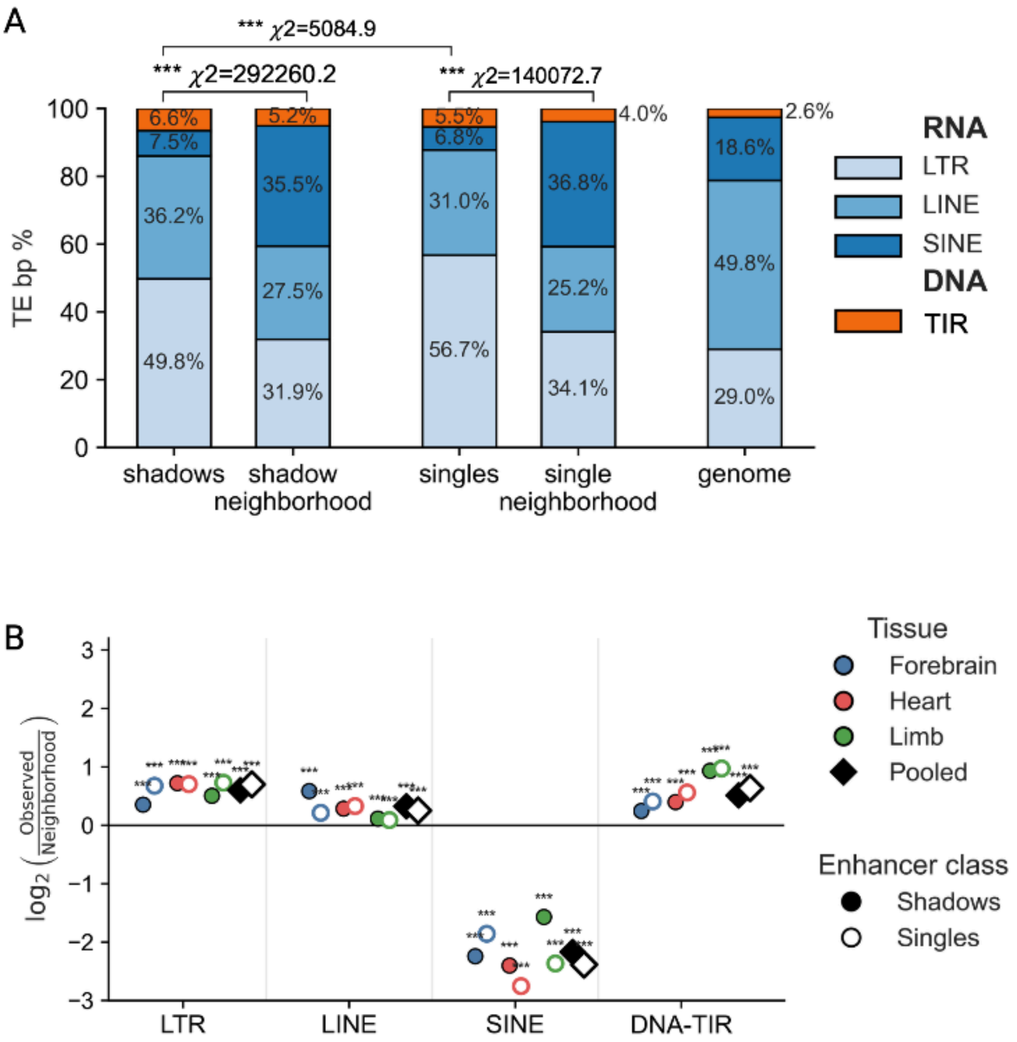


**Supplementary Figure 15. Mouse TE class composition among TE+ shadow and single enhancers, analyzed by basepair composition. (A)** Stacked bars show the base-pair composition of TE sequence overlapping TE co-opted shadow enhancers, TE+ single enhancers, their matched local neighborhoods, and the genome. This analysis is similar to Figure 6D and E, but analyzed by number of basepairs, instead of numbers of insertion events. Local neighborhoods were defined as the 50 kb upstream and downstream regions surrounding each TE+ enhancer. TE classes are grouped as RNA-derived elements, including LTR, LINE, and SINE elements, and DNA-derived elements, including TIR elements. Asterisks indicate significance from chi-square goodness-of-fit tests comparing observed TE class composition.**(B)** $Log_2$ enrichment of TE classes among TE+ enhancers relative to their matched local neighborhoods. Values show $log_2$(observed/neighborhood) TE base-pair frequency for shadow and single enhancers, shown separately by tissue and for the pooled dataset. Filled circles indicate shadow enhancers, open circles indicate single enhancers, and diamonds indicate pooled values. Asterisks indicate FDR-adjusted significance for per-class observed-versus-neighborhood enrichment tests.

**Supplementary Table 1. Fly BLAST metrics of enhancer-body hits of shadow pairs** (n=110 pair hits).

| BLAST Result Metrics | Value |
|---|---|
| % of shadow sets with hit | 14.6% |
| Median hit length | 18 bp |
| Range of hit length | [12 bps, 391 bps] |
| Median distance between shadow hit pair | 11,000 bps |
| Shadow hit distance range | [740 bps, 16,000 bps] |

**Supplementary Table 2. TE+ Shadow enhancer  fixation status of co-opted TEs across 10 *D. me*l lines.**

| Call category | n TEs | % of assessed (N=37) |
|---|---|---|
| Fixed (≥90% lines) | 27 | 72.97 |
| Not fixed (<90% lines) | 10 | 27.03 |
| Unassessed (call = NA) | 0 | — |

**Supplementary Table 3. TE+ Single enhancer fixation status of co-opted TEs across 10 *D. mel* lines.**

| Call category | n TEs | % of assessed (N=180) |
|---|---|---|
| Fixed (≥90% lines) | 169 | 93.89 |
| Not fixed (<90% lines) | 11 | 6.11 |
| Unassessed (call = NA) | 4 | — |

**Supplementary Table 4. Mouse BLAST metrics of enhancer-body hits of shadow pairs** (n=5,084 pair hits).

| BLAST Result Metrics | Value |
|---|---|
| % of shadow sets with hit | 31.9% |
| Median hit length | 87 bp |
| Range of hit length | [37 bp, 2,022 bp] |
| Median distance between shadow hit pair | 94,929 bp |
| Shadow hit distance range | [504 bp, 4,925,058 bp] |
